# Acetate enhances spatial memory in females via sex- and brain region-specific epigenetic and transcriptional remodeling

**DOI:** 10.1101/2024.08.26.609729

**Authors:** Erica M. Periandri, Kala M. Dodson, Francisca N. Vitorino, Benjamin A. Garcia, Karl M. Glastad, Gabor Egervari

## Abstract

Metabolic control of chromatin and gene expression is emerging as a key, but largely unexplored aspect of gene regulation. In the brain, metabolic-epigenetic interactions can influence critical neuronal functions. Here, we use a combination of behavioral, proteomic and genomic approaches to demonstrate that the intermediary metabolite acetate enhances memory in a brain region- and sex-specific manner. We show that acetate facilitates the formation of dorsal hippocampus-dependent spatial memories in female but not in male mice, while having no effect on cortex-dependent non-spatial memories in either sex. Acetate-enhanced spatial memory is driven by increased acetylation of histone variant H2A.Z, and upregulation of genes implicated in spatial learning in the dorsal hippocampus of female mice. In line with the sex-specific behavioral outcomes, the effect of acetate on dorsal hippocampal histone modifications and gene expression shows marked differences between the sexes during critical windows of memory formation (consolidation and recall). Overall, our findings elucidate a novel role for acetate, a ubiquitous and abundant metabolite, in regulating dorsal hippocampal chromatin, gene expression and learning, and outline acetate exposure as a promising new approach to enhance memory formation.

## INTRODUCTION

An emerging body of evidence suggests that epigenetic mechanisms are profoundly influenced by metabolic processes in the brain and other tissues (*1, 2*). Understanding how metabolism shapes chromatin and gene expression in the brain is critical, as gene regulatory mechanisms have been closely linked to key neuronal functions. Histone acetylation, for example, is strongly implicated both in adaptive processes, such as the formation of long-term memories (3, 4), and in the etiology of disease including substance use disorders (*5, 6*) or depression (7). A novel, unexpected mechanism of metabolic-epigenetic interactions is mediated by the nuclear translocation of metabolic enzymes such as ACSS2 (acetyl-CoA synthetase short chain family member 2). ACSS2 localizes to chromatin in differentiated neurons and regulates histone acetylation and gene expression by converting acetate into a local pool of acetyl-CoA, which is used by histone acetyltransferases to acetylate histones (8).

Recently, we showed that this pathway is critical for the incorporation of alcohol-derived acetate into histone acetylation in the brain, and that this incorporation drives the expression of genes related to learning and memory (9). We found that this epigenetic effect of alcohol-derived acetate is required for the formation of spatial memories in ethanol conditioned place preference. Consequently, inhibiting acetate-driven memory formation in the context of alcohol use could be a promising new therapeutic strategy (10). Given the key role of alcohol-derived acetate in facilitating alcohol-related memories, we hypothesize that the metabolite acetate could facilitate learning and memory formation in contexts independent of alcohol. If so, enhancing hippocampal memory formation via exposure to acetate might be of therapeutic relevance in conditions characterized by impaired cognitive performance. In further support, acetate induces transcriptional programs related to learning and memory in isolated primary hippocampal neurons (9). Whether the effects of acetate on learning and memory are generalizable to more adaptive contexts and different types of memory regulated by different brain regions, however, remains unknown.

To establish acetate as a memory enhancer, here we use a combination of behavioral, proteomic and genomic approaches. We show that acetate facilitates the formation of dorsal hippocampal spatial memories in female mice, while having no effect in males and not affecting cortex-dependent non-spatial learning. We find that acetate-enhanced spatial memory is accompanied by brain region- and sex-specific changes in histone modifications and gene expression both during consolidation and following recall. Our proteomic and transcriptomic data indicate that acetylation of the histone variant H2A.Z, a recently identified regulator of memory formation (11, 12), plays a key role in acetate-enhanced memory.

Overall, we establish the intermediary metabolite acetate as an important regulator of key epigenetic mechanisms and functionally relevant gene expression programs in the brain, and outline acetate exposure as a promising approach to facilitate dorsal hippocampal spatial memory formation in females. These results broaden our understanding of the role of metabolic-epigenetic interactions in the brain, and will drive the development of much-needed future therapies aiming to ameliorate memory impairments in aging, Alzheimer’s disease and other forms of dementia.

## RESULTS

### Acetate facilitates the formation of dorsal hippocampal spatial memory in female mice

To determine whether exogenous acetate facilitates learning in alcohol-independent contexts, we investigated its effect on two widely used behavioral assays of long-term memory in adult C57BL/6J mice (13). Given our previous findings showing the impact of alcohol-derived acetate on the dorsal hippocampus (dHPC) (9), we sought to examine whether acetate facilitates novel object location (NOL). NOL is an assay of dHPC-dependent spatial memory, in which mice learn to distinguish between familiar and novel locations of a specific object (13). Further, to determine whether acetate impacts other memory-related brain regions, we performed novel object recognition (NOR). NOR measures cortex-dependent non-spatial memory, in which mice differentiate between familiar and novel objects (13). Due to an innate preference of mice for novelty, animals with intact long-term memory tend to spend more time interacting with novel objects (NOR) or objects in a novel location (NOL) (13).

To test the effect of acetate on NOL and NOR behaviors, we exposed mice to 1.5 g/kg acetate or equivalent volume of saline, administered via intraperitoneal (i.p.) injection immediately prior to NOL or NOR training (***Fig. 1A***). Acetate-treated mice showed reduced distance traveled (***Fig. S1A,B***) and total object exploration time (***Fig. S1C,D***) during the NOL learning trial compared to saline-treated mice, suggesting that acetate exposure induces acute hypoactivity. Despite the decreased amount of time spent interacting with objects during the learning trial, acetate-treated mice tended to perform better during the NOL test trial (Student’s t test, t_36_=1.954, p=0.0585) (***Fig. 1B***): The average novelty preference index (NPI) of mice increased from 5.57 in the saline-treated group to 20.77 in the acetate-treated group.

**Figure 1.**
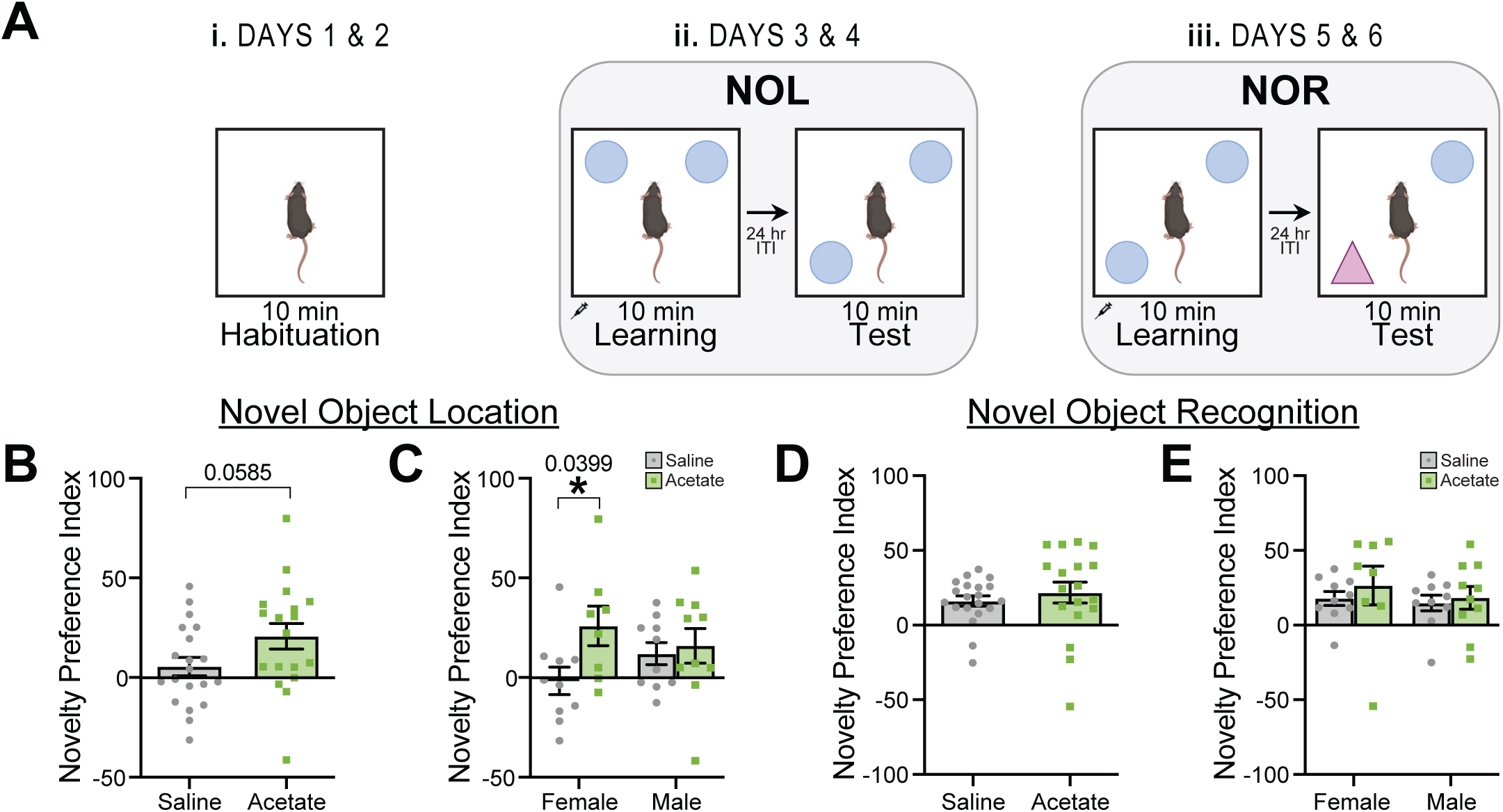
Acetate facilitates the formation of dorsal hippocampal spatial memory in female mice. **(A)** Schematic of behavioral battery. **(B)** Average novelty preference index (NPI) for the NOL test (Student’s t test t_36_=1.954, p=0.0585). **(C)** Average NPI for the NOL test, grouped by sex (two-way ANOVA interaction F_1,34_=2.203, p=0.1384; sex F_1,34_=0.05289, p=0.8195; treatment F_1,34_=4.120, p=0.0503; post-hoc Šídák’s multiple comparisons t_34_=2.438, p-adj.=0.0399 for females and t_34_=0.3833, p-adj.=0.9166 for males). **(D)** Average NPI for the NOR test (Student’s t test t_36_=0.7437, p=0.4619). **(E)** Average NPI for the NOR test, grouped by sex (two-way ANOVA interaction F_1,34_=0.1128, p=0.7391; sex F_1,34_=0.5381, p=0.4683; treatment F_1,34_=0.6183, p=0.4371; post-hoc Šídák’s multiple comparisons t_34_=0.9118, p-adj.=0.6930 for females and t_34_=0.2893, p-adj.=0.9348 for males). Sample sizes are n_Saline,F_=10, n_Saline,M_=10, n_Acetate,F_=8, n_Acetate,M_=10. Abbreviations: NOL, novel object location; NOR, novel object recognition; NPI, novelty preference index.

Previous studies have reported sex differences in the behavioral responses to both spatial and object novelty in adult C57BL/6J mice (14). Indeed, when stratifying NPI by sex, we found that the facilitating effect of acetate on NOL was driven by females (***Fig. 1C***). In female mice, acetate led to a significant increase of NPI from an average of -1.22 in the saline group to an average of 23.66 in the acetate treated group (post-hoc Šídák’s test, t_34_=2.438, p-adj.=0.0399), while no effect was found in males (post-hoc Šídák’s test, t_34_=0.3733, p-adj.=0.9166).

Similarly to NOL, acetate-treated mice exhibited acute hypoactivity during the NOR learning trial, as evidenced by significantly reduced distance traveled and a trend towards decreased total interaction time (***Fig. S1E-H***). In contrast with NOL, however, acetate had no effect on NPI during NOR when combining females and males (Student’s t test, t_36_=0.7437, p=0.4619) (***Fig. 1D***), nor when stratified by sex (post-hoc Šídák’s test, t_34_=0.7711, p-adj.=0.6930 in females and t_34_=0.3284, p-adj.=0.9348 in males) (***Fig. 1E***).

These results indicate that acetate treatment enhances dHPC-dependent spatial memory in female but not in male mice, while having no effect on cortex-dependent non-spatial learning in either sex. Taken together, our findings suggest that exogenous acetate facilitates memory formation in a sex- and brain region-specific manner.

### Increased acetylation of histone variant H2A.Z in the dorsal hippocampus of acetate-treated female mice

We next sought to elucidate the molecular underpinnings of acetate-enhanced spatial learning. We previously reported that alcohol-derived acetate fuels dorsal hippocampal histone acetylation in the context of alcohol-associated learning (9). Therefore, we hypothesized that acetate might also lead to changes in histone post-translational modifications (hPTMs) in the context of dHPC-dependent memory formation. To characterize hPTMs in the brains of acetate- or saline-treated mice that underwent NOL and NOR, we performed liquid chromatography coupled with tandem mass spectrometry (LC-MS/MS) targeting histones. Given the brain region and sex specificity of the behavioral outcomes, we tested both dHPC and cortex in both male and female mice following memory recall.

In line with the behavioral effects described in ***Fig. 1***, we found that acetate impacted chromatin modifications in both a sex and brain region-specific manner. In females, we found that acetate treatment significantly changed a larger number of hPTMs in the dHPC (n=27 histone peptides with p<0.05) compared to the cortex (n=9 histone peptides with p<0.05) (***Fig. 2A***). In contrast, the cortex was more affected (n=23 histone peptides with p<0.05) than the dHPC (n=7 histone peptides with p<0.05) in male mice (***Fig. 2B***). Further emphasizing the different responses to acetate between sexes, the dHPC had few significantly changed peptides when considering females and males together (***Fig. S2A***).

**Figure 2.**
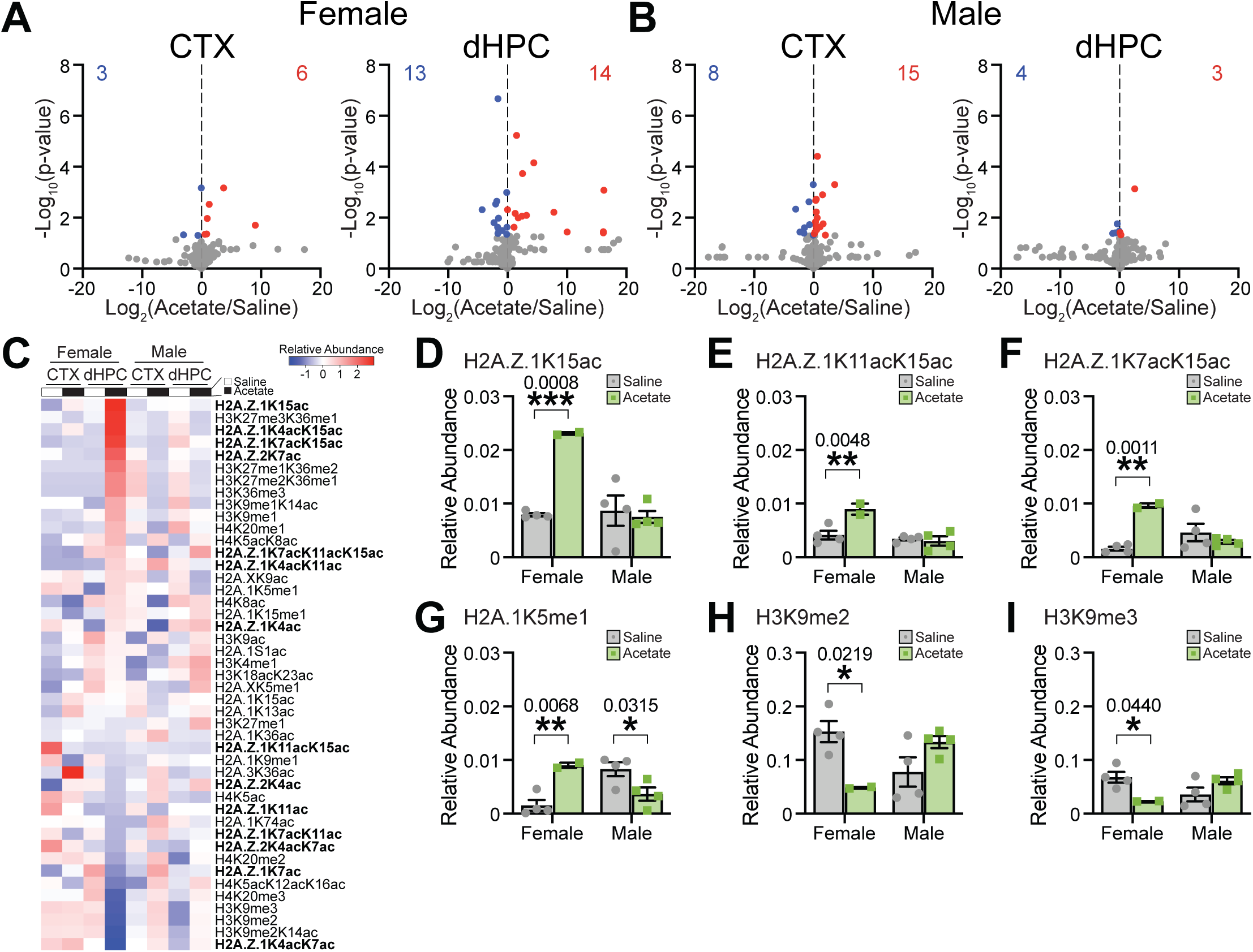
Acetate increases the acetylation of histone variant H2A.Z in the female dorsal hippocampus. **(A)** Volcano plots of hPTMs in the CTX (left) and dHPC (right) comparing acetate- and saline-treated female mice following memory recall (late timepoint). **(B)** Volcano plots of hPTMs in the CTX and dHPC comparing acetate- and saline-treated male mice sacrificed following memory recall. **(C)** Heatmap of average row Z-score of relative abundance of significantly affected hPTMs, sorted from highest to lowest in the acetate-treated female dHPC. H2A.Z hPTMs are bolded. **(D-I)** Average relative abundance of hPTMs of interest in the dHPC following memory recall. **(D)** Average relative abundance of H2A.Z.1K15ac (two-way ANOVA interaction F_1,10_=19.01, p=0.0014; post-hoc Šídák’s multiple comparisons t_10_=5.224, p-adj.=0.0008 for females and t_10_=0.4963, p-adj.=0.8634 for males). **(E)** Average relative abundance of H2A.Z.1K11acK15ac (two-way ANOVA interaction F_1,10_=11.39, p=0.0071; post-hoc Šídák’s multiple comparisons t_10_=4.031, p-adj.=0.0048 for females and t_10_=0.3981, p-adj.=0.9094 for males). **(F)** Average relative abundance of H2A.Z.1K7acK15ac (two-way ANOVA interaction F_1,10_=21.89, p=0.0009; post-hoc Šídák’s multiple comparisons t_10_=4.966, p-adj.=0.0011 for females and t_10_=1.317, p-adj.=0.3875 for males). **(G)** Average relative abundance of H2A.1K5me1 (two-way ANOVA interaction F_1,10_=22.89, p=0.0007; post-hoc Šídák’s multiple comparisons t_10_=3.810, p-adj.=0.0068 for females and t_10_=2.898, p-adj.=0.0315 for males). **(H)** Average relative abundance of H3K9me2 (two-way ANOVA interaction F_1,10_=13.70, p=0.0041; post-hoc Šídák’s multiple comparisons t_10_=3.113, p-adj.=0.0219 for females and t_10_=2.040, p-adj.=0.1326 for males). **(I)** Average relative abundance of H3K9me3 (two-way ANOVA interaction F_1,10_=10.84, p=0.0081; post-hoc Šídák’s multiple comparisons t_10_=2.702, p-adj.=0.0440 for females and t_10_=1.896, p-adj.=0.1667 for males). For volcano plots, dots represent individual hPTMs (gray not significant, blue significantly downregulated, and red significantly upregulated by acetate vs. saline), and numbers denote the quantity of significantly downregulated (blue, top left) or significantly upregulated (red, top right) hPTMs. *p<0.05, **p<0.01, ***p<0.001. Sample sizes are n_Saline,F_=4, n_Saline,M_=4, n_Acetate,F_=2, n_Acetate,M_=4. Abbreviations: CTX, cortex; dHPC, dorsal hippocampus; hPTMs, histone post-translational modifications.

Looking at individual hPTMs, we found that over 30% of all significantly affected histone modifications were acetylation marks on the histone variant H2A.Z (***Fig. 2C***), which has been previously linked to long-term memory (12). In the dHPC, this effect was remarkably sex-specific: while H2A.Z acetylation marks comprised over 40% of upregulated and 15% of downregulated hPTMs in females, no changes of H2A.Z acetylation were observed in the males. Further, the induction of H2A.Z acetylation was highly consistent across various histone peptides detected in LC-MS/MS. In the female dHPC, we found a significant increase in the average relative abundance of H2A.Z.1K15ac (two-way ANOVA interaction F_1,10_=19.01, p=0.0014; post-hoc Šídák’s multiple comparisons test t_10_=5.224, p-adj.=0.0008 in females and t_10_=0.4963, p-adj.=0.8634 in males) (***Fig. 2D***), H2A.Z.1K11acK15ac (two-way ANOVA interaction F_1,10_=11.39, p=0.0071; post-hoc Šídák’s multiple comparisons test t_10_=4.031, p-adj.=0.0048 in females and t_10_=0.3981, p-adj.=0.9094 in males) (***Fig. 2E***), and H2A.Z.1K7acK15ac (two-way ANOVA interaction F_1,10_=21.89, p=0.0009; post-hoc Šídák’s multiple comparisons test t_10_=4.966, p-adj.=0.0011 in females and t_10_=1.317, p-adj.=0.3875 in males) (***Fig. 2F***). None of these peptides were significantly induced by acetate in the male dHPC, nor in the cortex of either sex (***Fig. S2B-D***).

To corroborate these findings, we performed Western blot analysis of H2A.Z pan-acetylation levels in the dHPC of female mice 24 hours following i.p. injection of 1.5 g/kg acetate or equivalent volume of saline. We found increased H2A.Z acetylation in the dHPC of female mice following acetate exposure (***Fig. S2H***), supporting the significantly increased dorsal hippocampal H2A.Z acetylation in acetate-treated females observed in LC-MS/MS.

In addition to H2A.Z, acetate also affected hPTMs on other histone H2A peptides, such as H2A.1 (***Fig. 2C***). For example, we found a sex-specific effect of acetate on H2A.1K5me1 in the dHPC (interaction from two-way ANOVA F_1,10_=22.89, p=0.0007), with increased mono-methylation in females (post-hoc Šídák’s multiple comparisons test t_10_=3.81, p-adj.=0.0068) and decreased mono-methylation in males (post-hoc Šídák’s multiple comparisons test t_10_=2.898, p-adj.=0.0315) (***Fig. 2G***). No changes of H2A.1 K5me1 were observed in the cortex (***Fig. S2E***).

Finally, we found significantly decreased methylation of repressive mark H3K9 in the dHPC acetate-treated females. This effect was consistent both for H3K9me2 (two-way ANOVA interaction F_1,10_=13.70, p=0.0041; post-hoc Šídák’s multiple comparisons test t_10_=3.113, p-adj.=0.0219 in females and t_10_=2.04, p-adj.=0.1326 in males) (***Fig. 2H***) and H3K9me3 (two-way ANOVA interaction F_1,10_=10.84, p=0.0081; post-hoc Šídák’s multiple comparisons test t_10_=2.702, p-adj.=0.0440 in females and t_10_=1.896, p-adj.=0.1667 in males) (***Fig. 3I***). No changes of H3K9me were observed in the cortex (***Fig. S2F,G***).

**Figure 3.**
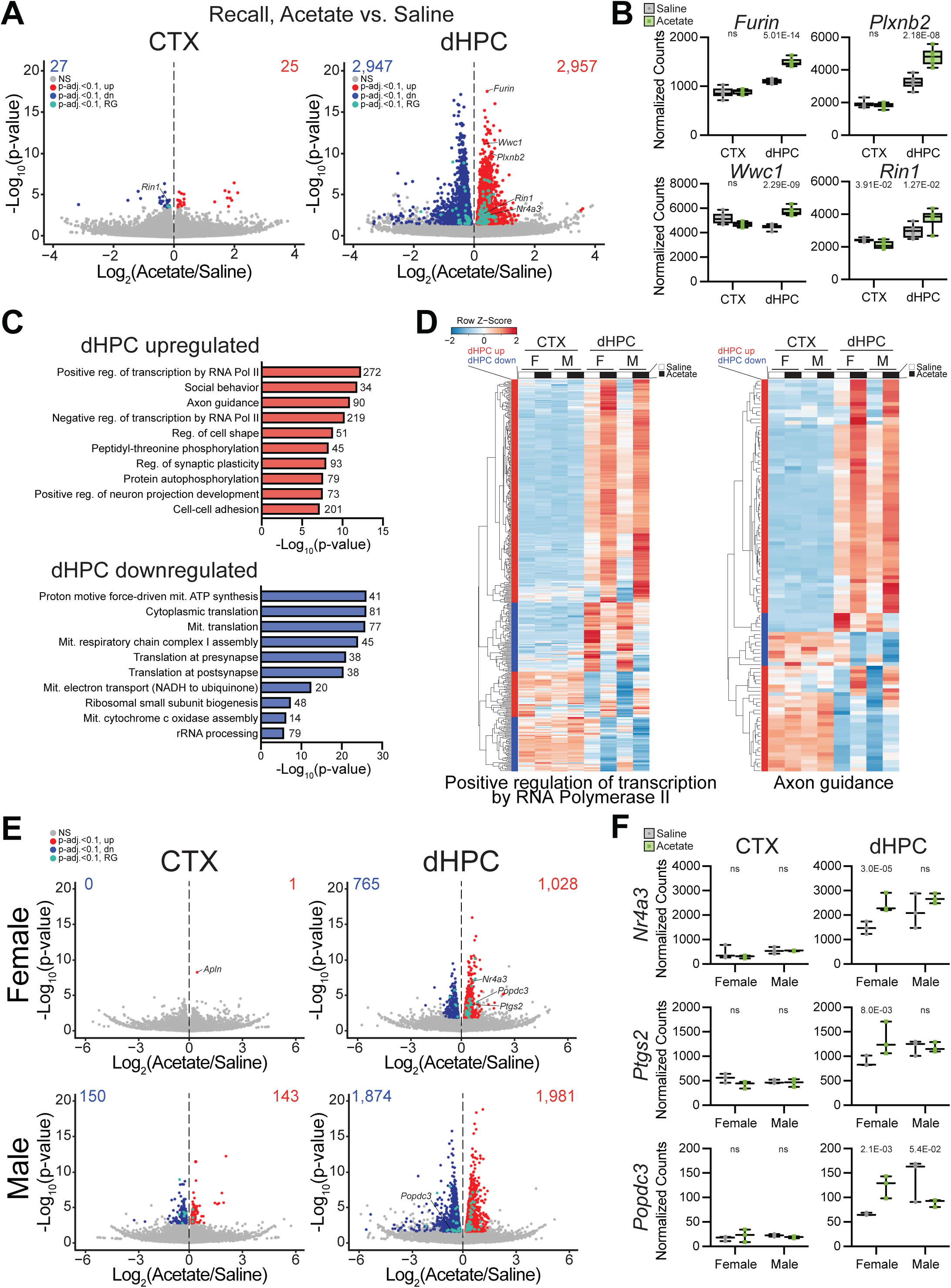
Acetate enhances the expression of learning and memory-related genes in the female dorsal hippocampus. **(A)** Volcano plots showing differential gene expression in the CTX (left) and dHPC (right) of acetate-vs. saline-treated mice following memory recall (late timepoint). **(B)** Normalized gene counts for GOIs *Furin* (top left), *Plxnb2* (top right), *Wwc1* (bottom left), and *Rin1* (bottom right). **(C)** Top 10 significantly enriched GO terms in the dHPC for upregulated DEGs (top) and downregulated DEGs (bottom). Numbers next to bars indicate the quantity of significant DEGs within each corresponding GO term. **(D)** Heatmaps of average row Z-score of TxPM for all dHPC DEGs associated with two significantly enriched GO terms of interest: positive regulation of transcription by RNA Polymerase II (left) and axon guidance (right). The colored bar between each dendrogram and heatmap indicates directionality of differential expression comparing acetate-vs. saline-treated dHPC leveraging both sexes (red upregulated, blue downregulated). **(E)** Volcano plots showing differential gene expression in the CTX (left) and dHPC (right) of acetate-vs. saline-treated females (top) and males (bottom) following memory recall. **(F)** Normalized gene counts for response GOIs *Nr4a3* (top), *Ptgs2* (middle), and *Popdc3* (bottom) in CTX (left) and dHPC (right). For volcano plots, dots represent individual genes; gray represents not significant (NS), red significantly upregulated by acetate, blue significantly downregulated by acetate, teal significant and response gene (RG). Numbers denote the quantity of significantly downregulated (blue, top left) or significantly upregulated (red, top right) genes. Sample sizes are n=3 per sex per condition. Abbreviations: CTX, cortex; dHPC, dorsal hippocampus; F, female; GO, gene ontology; GOI, gene of interest; M, male; mit., mitochondrial; Nr4a3, nuclear receptor subfamily 4, group A, member 3; NS, not significant; Plxnb2, plexin B2; Popdc3, popeye domain containing 3; Ptgs2, prostaglandin-endoperoxide synthase 2; Rin1, Ras and Rab interactor 1; RG, response gene; reg., regulation; TxPM, transcripts per million; Wwc1, WW, C2, and coiled-coil domain containing 1.

Taken together, our LC-MS/MS data indicate that acetate has a sex- and brain-region specific effect on histone post-translational modifications in the context of learning. This included a remarkably consistent upregulation of H2A.Z acetylation in the female dHPC, not otherwise observed in the male dHPC nor in the cortex of either sex. Dorsal hippocampal H2A.Z acetylation has been previously linked to spatial memory (12), further suggesting that this histone variant might play a critical role in acetate-enhanced spatial learning in females.

### Acetate enhances the expression of genes related to learning and memory in the female dorsal hippocampus

We next profiled the effect of acetate on gene expression changes in the dHPC and cortex using RNA sequencing. Emphasizing regional specificity, we identified several differentially expressed genes (DEGs) associated with NOL learning performance in the dHPC of saline-injected control mice (143 upregulated and 2 downregulated genes when comparing mice with positive NPI to mice with negative NPI) but not in the cortex (9 upregulated and 8 downregulated genes). Likewise, gene expression was associated with NOR learning performance in the cortex (28 upregulated and 7 downregulated DEGs) but not in the dHPC (0 upregulated and 2 downregulated DEGs) (***Fig. S3***). This marked dichotomy is in line with the critical role of the dHPC and cortex in regulating spatial and non-spatial memory formation, respectively, and suggests that our transcriptional profiling captured behaviorally relevant gene expression states.

In line with the strong effect of acetate on dorsal hippocampal histone modifications and spatial memory, we found that acetate treatment resulted in a considerably more robust dysregulation of gene expression in the dHPC (n=5,904 genes) compared to the cortex (n=52 genes) (***Fig. 3A***). Several DEGs that were upregulated in the dHPC of acetate-treated mice have been implicated in learning and memory, including the most significantly upregulated gene *Furin* (furin, paired basic amino acid cleaving enzyme), as well as *Plxnb2* (plexin B2), *Wwc1* (WW, C2, and coiled-coil domain containing 1), and *Rin1* (Ras and Rab interactor 1) (***Fig. 3B***) (15–18). None of these genes were significantly upregulated by acetate in the cortex; in fact, *Rin1* expression was significantly decreased in this brain region (***Fig. 3B*, *bottom right***). Of note, acetate treatment resulted in a predominant upregulation of response genes in the dHPC (***Fig. 3A, teal dots***), including *Nr4a3*, *Kcnj4*, *Nptx2*, *Gpr63*, *Kcna1*, *Egr4* and *Hunk*. Response genes play a critical role in orchestrating activity-dependent changes in neurons, and have been tightly linked to spatial memory formation in the dHPC (19).

Gene ontology (GO) analysis of significantly upregulated dorsal hippocampal genes revealed an enrichment of genes related to both positive and negative regulation of transcription by RNA Polymerase II, as well as axon guidance and regulation of synaptic plasticity (***Fig. 3C, top***), suggesting that the acetate-induced changes in hippocampal gene expression play an important role in the observed learning phenotype (***Fig. 1***). Most genes in these categories were significantly increased by acetate (***Fig. 3D***), which is concordant with the strong induction of histone acetylation in the dHPC. In contrast, GO analyses of downregulated dorsal hippocampal genes revealed enrichment of genes related to translation and mitochondrial processes (***Fig. 3C, bottom***), possibly as a compensatory effect to the metabolic perturbations caused by exogenous acetate.

The larger effect of acetate on dorsal hippocampal transcription was evident in both sexes, but particularly in females, where only 1 gene was differentially expressed in the cortex, compared to 1,793 in the dHPC (***Fig. 3E***). A closer look at the expression pattern of response genes revealed intriguing differences between the brain regions and sexes. In the dHPC, the effect of acetate on response genes showed a clear sex-specific difference with predominant upregulation in females (31 genes significantly increased and 5 genes significantly decreased), and no directional bias in males (37 genes significantly increased and 27 genes significantly decreased) (***Fig. 3E***). For example, we found a significant increase of *Nr4a3* (Nuclear receptor subfamily 4 group a member 3) expression in the dHPC of acetate-treated females, whereas no changes were observed in the male dHPC nor in the cortex of either sex (***Fig. 3F, top***). *Nr4a3* is a primary response gene encoding for a nuclear receptor transcription factor, which is a critical regulator of transcription-dependent memory consolidation and has been proposed as a target to improve cognitive functions (20, 21). Similarly, *Ptgs2* (prostaglandin-endoperoxidase synthase 2) is a response gene that was significantly increased in the dHPC of acetate-treated females (***Fig. 3F, middle***), but was not affected in males nor in the cortex. The expression of response gene *Popdc3* (Popeye domain-containing protein 3) was significantly increased by acetate in the dHPC in females, and significantly decreased in the dHPC of males (***Fig. 3F, bottom***), with no changes in the cortex of either sex.

Overall, our transcriptional profiling revealed that acetate significantly upregulates the expression of genes previously implicated in learning and memory. Learning-related gene expression changes were most pronounced in the female dHPC, which was in line with the sex- and brain region-specific effects of acetate on histone modifications and learning behavior.

### Acetate regulates the dorsal hippocampal expression of genes linked to novel object location memory in a sex-specific manner

To further characterize the potential role of acetate-induced transcriptional changes in enhancing spatial memory in female mice, we compared genes regulated by acetate to genes that were differentially expressed between mice with positive NPIs (learners) and mice with negative NPIs (non-learners) in NOL (e.g., ***Fig. S3A***). This analysis revealed a clear directional overlap in which acetate-upregulated genes were significantly enriched among NOL-upregulated genes (Fisher’s exact test, p=4.2E-04) and acetate-downregulated genes were significantly enriched among NOL-downregulated genes (Fisher’s exact test, p=9E-08) (***Fig. 4A***). These effects were driven by females, with highly significant overlaps between both acetate- and NOL-upregulated genes (Fisher’s exact test, p=7.6E-12) and acetate- and NOL-downregulated genes (Fisher’s exact test, p=5.1E-08) (***Fig. 4B, left***). In males, acetate-upregulated genes showed no significant overlap with either NOL-upregulated or NOL-downregulated genes and, surprisingly, acetate-downregulated genes were enriched among NOL-upregulated genes (Fisher’s exact test, p=4.3E-11) (***Fig. 4B, right***). Overall, these analyses indicate that acetate induced the expression of genes associated with NOL learning specifically in the female dHPC, which is in line with the observed sex-specific differences in facilitating spatial memory.

**Figure 4.**
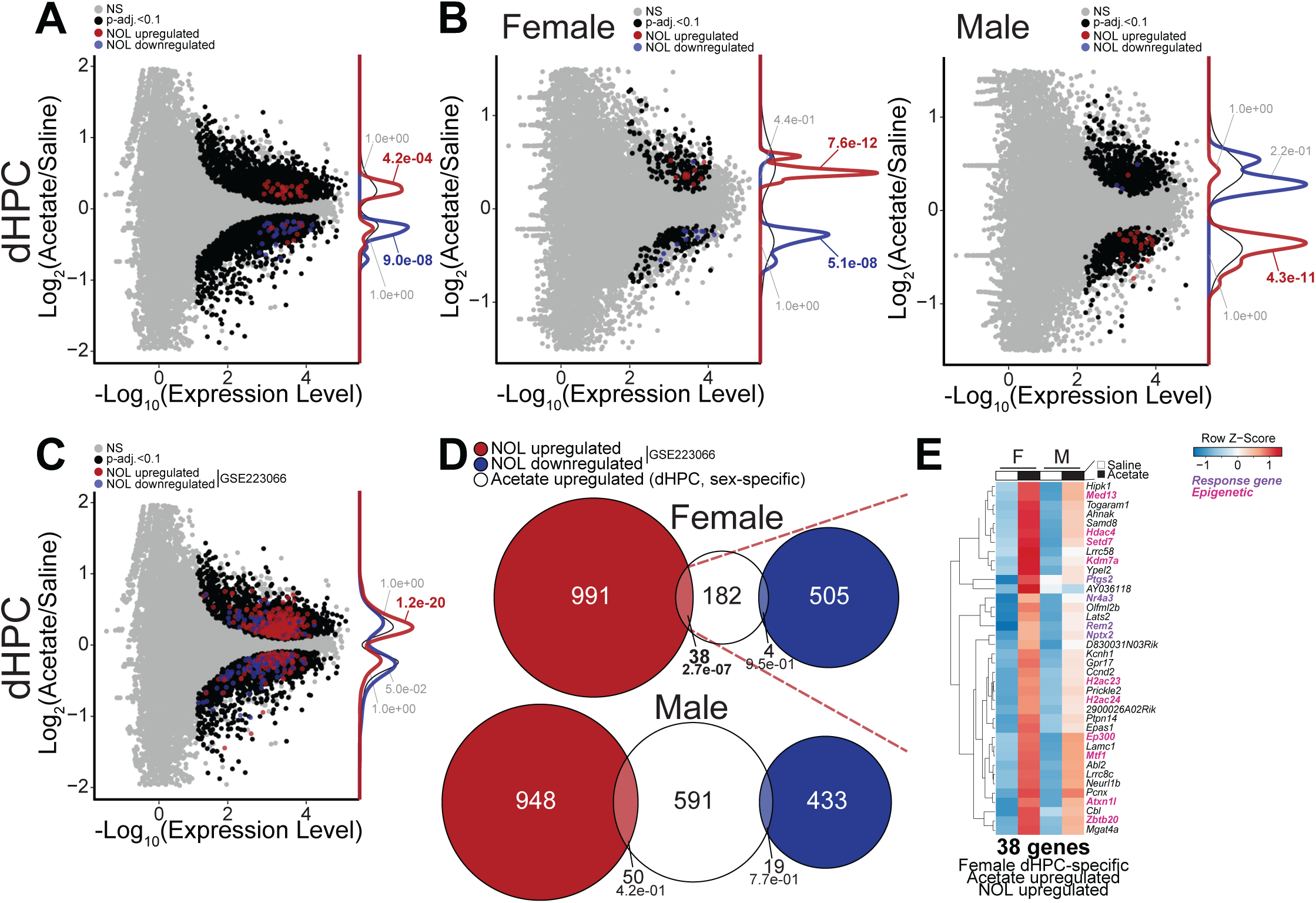
Acetate increases the dorsal hippocampal expression of novel object location memory-linked genes in a sex-specific manner. **(A)** MA plot showing the relationship of gene expression changes in response to acetate treatment and learning in the NOL task in the dHPC. Overlaps were analyzed using Fisher’s exact test (corresponding p-values displayed). Density plots of associated overlapping gene categories and significance of overlap (Fisher’s exact test) given to the right of plot. **(B)** MA plots showing the relationship of gene expression changes in response to acetate treatment and NOL learning in the dHPC of females (left) and males (right). **(C)** MA plot showing the relationship of gene expression changes in response to acetate treatment and NOL learning (DEGs from (22)) in the dHPC. **(D)** Venn diagrams showing overlap of DEGs in response to acetate treatment (acetate upregulated, white) and NOL learning (from (22); NOL upregulated red, NOL downregulated blue) in the dHPC of females (top) and males (bottom). Overlaps were analyzed using Fisher’s exact test (corresponding p-values and number of DEGs displayed). **(E)** Heatmap of average row Z-score of TxPM for 38 DEGs upregulated in response to acetate treatment and NOL learning, identified by (22) in the female dHPC. Purple gene names indicate response genes, and pink gene names indicate genes with epigenetic function. For MA plots, dots represent individual transcripts; gray represents not significant (NS), black significant (p-adj<0.1), red significant and upregulated in NOL learners (NOL upregulated), blue significant and downregulated in NOL learners (NOL downregulated). NOL learners are characterized by a positive NPI for the NOL test, whereas non-learners have a negative NPI for the NOL test. Histograms show the distribution of DEGs overlapping between acetate treatment and NOL learning. Sample sizes are n=3 per sex per group. Abbreviations: CTX, cortex; DEGs, differentially expressed genes; dHPC, dorsal hippocampus; F, female; M, male; MA, mean average; NOL, novel object location; TxPM, transcripts per million.

To corroborate these findings, we next compared our gene expression data to a previously published study in which RNA-seq of the dHPC was performed in the context of NOL (22). Once again, we found a strong directional overlap between genes affected by acetate in the dHPC and genes that were changed in the dHPC following NOL (***Fig. 4C***). This was particularly evident for genes upregulated by both acetate and NOL (Fisher’s exact test, p=1.2E-20), and also reached significance for genes downregulated by acetate and NOL (Fisher’s exact test, p=0.048). Of note, this directional overlap was primarily driven by females. Specifically, genes upregulated by acetate in the female dHPC showed a highly significant overlap with NOL upregulated genes (Fisher’s exact test, p=2.7E-07) but not with NOL downregulated genes (Fisher’s exact test, p=0.95) (***Fig. 4D***). In contrast, genes upregulated by acetate in the male dHPC showed no significant overlaps with either NOL upregulated (Fisher’s exact test, p=0.42) or NOL downregulated (Fisher’s exact test, p=0.77) genes (***Fig. 4D***).

Genes that were upregulated by both acetate and NOL in the female dHPC showed a more subtle increase in the male dHPC (***Fig. 4E***), emphasizing the sex-specific effects of acetate. Over 25% of these genes had chromatin-related functions (***Fig. 4E***), including *Hdac4*, *Hdac23*, and *Hdac24*, which suggests a potential compensatory increase of histone deacetylase enzymes to counteract the acetate-induced increase in dorsal hippocampal histone acetylation. Additional acetate- and NOL-upregulated genes included histone acetyltransferase *Ep300*, histone methyltransferase *Setd7* and histone demethylase *Kdm7a*, as well as several response genes including *Nr4a3*, *Nptx2* and *Ptgs2* (***Fig. 4E***).

Taken together, we found that acetate increased the expression of genes previously linked to NOL in the female dHPC, further emphasizing the important role of acetate-driven transcriptional changes in facilitating spatial memory formation in female mice.

### Genes upregulated by acetate in the female dorsal hippocampus are linked to histone variant H2A.Z

Considering the sex- and brain-region specific hyperacetylation of H2A.Z in acetate-treated mice (***Fig. 2***), we next explored the relationship between acetate-affected genes and genes regulated by H2A.Z. First, we tested the overlap between our DEGs and genes that were downregulated in the hippocampus following cre-dependent knock-out (KO) of H2A.Z expression (12). We found that both acetate-upregulated (Fisher’s exact test, p=7.0E-25) and acetate-downregulated (Fisher’s exact test, p=5.1E-14) genes significantly overlapped with genes that were downregulated by H2A.Z KO, suggesting an important role for H2A.Z in mediating acetate-related transcriptional changes (***Fig. 5A***). After stratifying data by sex, we found the most significant overlap in female mice between genes upregulated by acetate and genes downregulated by H2A.Z KO (Fisher’s exact test, p=3.3E-06) (***Fig. 5B***). Further, genes downregulated by acetate also showed a significant overlap with genes downregulated by H2A.Z KO (Fisher’s exact test, p=6.7E-03) in females. In stark contrast, genes downregulated by H2A.Z KO showed no significant overlaps with acetate-upregulated (Fisher’s exact test, p=0.97) nor acetate-downregulated (Fisher’s exact test, p=0.38) genes in the dHPC of male mice (***Fig. 5B***).

**Figure 5.**
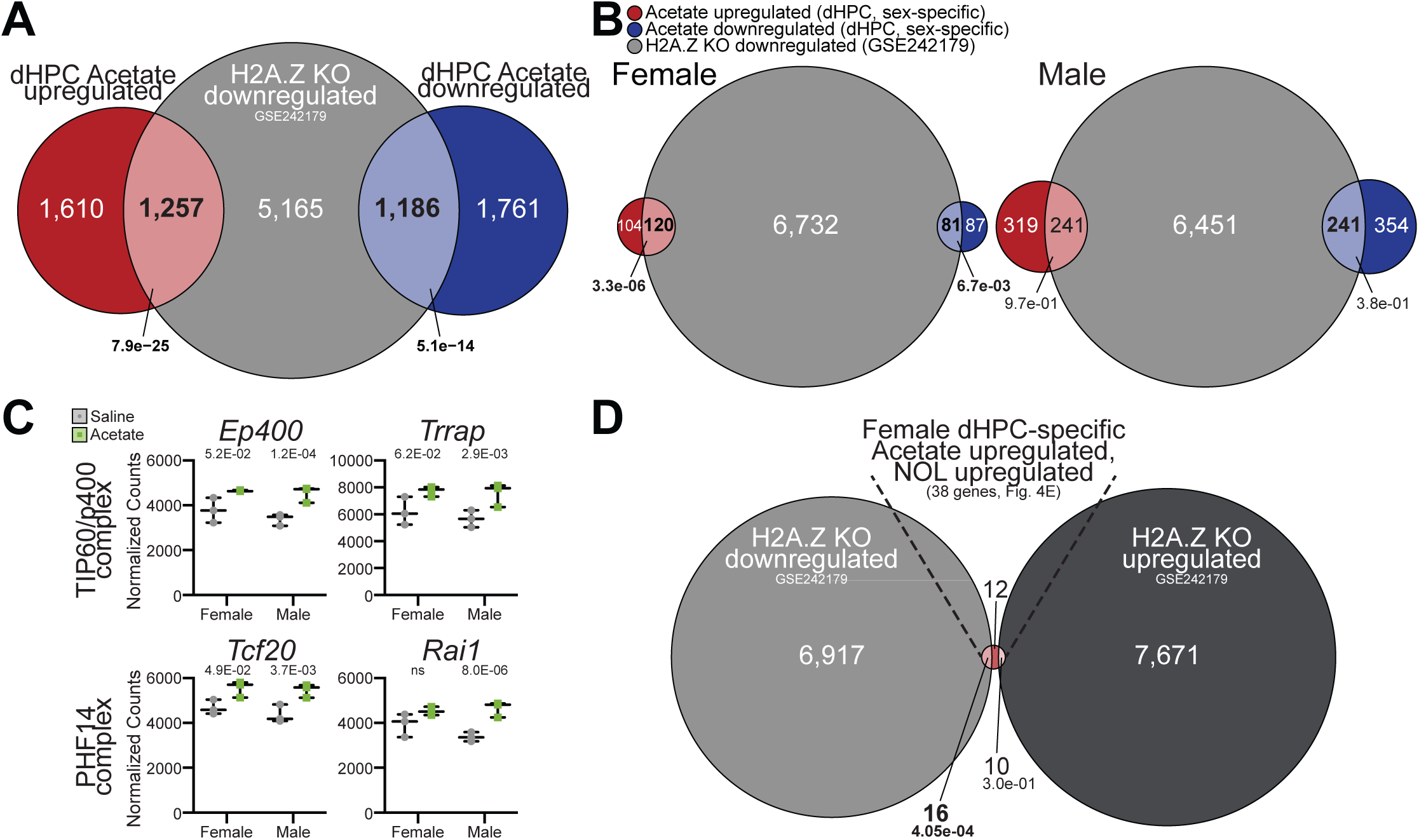
Genes upregulated by acetate in the female dorsal hippocampus are linked to histone variant H2A.Z. **(A)** Venn diagram showing overlap of DEGs in the dHPC in response to acetate treatment following memory recall (late timepoint) and deletion of H2A.Z, identified by (12). **(B)** Venn diagrams showing overlap of DEGs in the dHPC in response to acetate treatment following memory recall (acetate upregulated red, acetate downregulated blue) and deletion of H2A.Z, identified by (12) in females (left) and males (right). **(C)** Normalized gene counts in the dHPC for GOIs associated with H2A.Z deposition and acetylation, identified by (23). *Ep400* and *Trrap* encode components of the TIP60/p400 complex, and *Tcf20* and *Rai1* encode components of the PHF14 complex. P-values represent DESeq2 adjusted p-values. **(D)** Venn diagram showing overlap of DEGs in the dHPC in response to deletion of H2A.Z (downregulated, gray and upregulated, dark gray) and the 38 female DEGs upregulated in the context of acetate and NOL learning (identified in Fig. 4E). For Venn diagrams, overlaps were analyzed using Fisher’s exact test (corresponding p-values and number of DEGs displayed). Sample sizes are n=3 per sex per group. Abbreviations: DEGs, differentially expressed genes; dHPC, dorsal hippocampus; Ep400, E1A binding protein p400; KO, knockout; ns, not significant; PHF14, PHD finger protein 14; Rai1, retinoic acid induced 1; Tcf20, transcription factor 20; TIP60 (KAT5), lysine acetyltransferase 5; Trrap, transformation/transcription domain-associated protein.

We next examined how acetate affects the expression of known H2A.Z chaperones and acetyltransferases (23). We found that acetate increased the expression of *Ep400* and *Trrap* (***Fig. 5C***), two members of the TIP60/p400 complex, which has been implicated in both H2A.Z deposition and acetylation (24). In addition, acetate also increased the expression of *Tcf20* and *Rai1* (***Fig. 5C***), members of the PHF14 complex, which mediates the effect of H2A.Z on gene expression (23). Acetate did not affect the expression of any of these genes in the cortex of either sex (***Fig S3D***).

Of the 38 genes that were induced by both acetate and NOL in the dHPC of female mice, 16 were decreased by H2A.Z deletion (Fisher’s exact test, p=4E-04), whereas the overlap with genes upregulated by H2A.Z deletion was not significant (Fisher’s exact test, p=0.3) (***Fig. 5D***). This directional overlap suggests that the upregulation of acetate- and NOL-induced genes depends on H2A.Z expression.

Taken together, these results suggest that the histone variant H2A.Z plays an important role in mediating acetate-induced gene expression changes in the dorsal hippocampus, further supporting our mass spectrometry findings.

### Acetate induces sex-specific changes of histone modifications and gene expression in the dorsal hippocampus during memory consolidation

Our main analysis focused on the molecular landscape of the dHPC and cortex following the establishment of long-term memories, approximately 24 hours after acetate exposure and immediately following memory recall. We next sought to examine hPTMs and gene expression in the d HPC of mice during the memory consolidation period. To this end, a second cohort of mice received i.p. injections of 1.5 g/kg acetate or equivalent volume of saline prior to NOL training, and was sacrificed immediately following the learning session.

First, we performed LC-MS/MS to characterize hPTMs in the dHPC. When combining sexes, acetate treatment resulted in 14 hPTM changes during the consolidation window (***Fig. S4A***). In clear contrast with the later time point, there were fewer chromatin changes in the dHPC of females (n=7 upregulated peptides and n=1 downregulated peptide) compared to males (n=14 upregulated peptides and n=5 downregulated peptides) (***Fig. 6A***). Acetate had a more pronounced effect on canonical histones compared to histone variants at this early time point (***Fig. 6B***). For example, the relative abundance of double acetylated H4K8acK12ac significantly increased in females but not in males (Šídák’s multiple comparisons t_8_=5.215, p-adj.=0.0016 for females and t_8_=0.6114, p-adj.=0.8045 for males) (***Fig. 6C***). In contrast, the relative abundance of quadruple acetylated H4K5acK8acK12acK16ac did not change in females but significantly increased in males (Šídák’s multiple comparisons t_8_=0.0741, p-adj.=0.9967 for females and t_8_=4.144, p-adj.=0.0065 for males) (***Fig. 6D***). Similarly, the relative abundance of double acetylated H3K18acK23ac and H2A.Z.1 K4acK7ac did not change in females but significantly increased in males (Šídák’s multiple comparisons t_8_=1.444, p-adj.=0.3384 for females and t_8_=3.279, p-adj.=0.0223 for males; t_8_=1.342, p-adj.=0.3862 for females and t_8_=4.052, p-adj.=0.0073 for males) (***Fig. 6E,F***).

**Figure 6.**
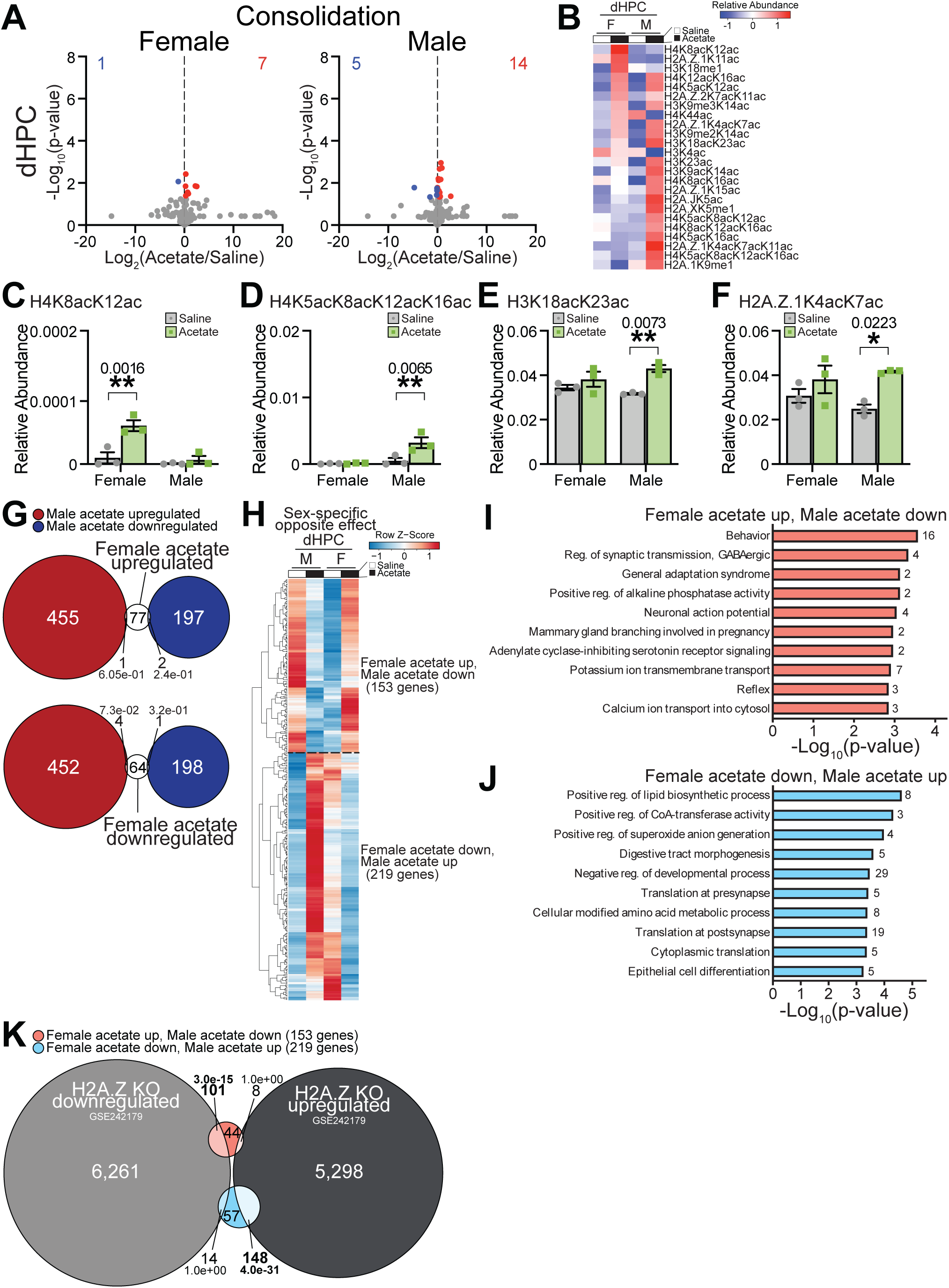
Acetate induces sex-specific changes of histone modifications and gene expression in the dorsal hippocampus during memory consolidation. **(A)** Volcano plots of hPTMs in the dHPC comparing acetate- and saline-treated female (left) and male (right) mice sacrificed following the NOL learning trial, during consolidation (early timepoint). **(B)** Heatmap of average row Z-score of relative abundance of significant hPTMs, sorted from highest to lowest in the acetate-treated female dHPC. **(C-F)** Average relative abundance of hPTMs of interest in the dHPC during memory consolidation. **(C)** Average relative abundance of H4K8acK12ac (two-way ANOVA interaction F_1,8_=10.60, p=0.0116; post-hoc Šídák’s multiple comparisons t_8_=5.215, p-adj.=0.0016 for females and t_8_=0.6114, p-adj.=0.8045 for males). **(D)** Average relative abundance of H4K5acK8acK12acK16ac (two-way ANOVA interaction F_1,8_=8.282, p=0.0206; post-hoc Šídák’s multiple comparisons t_8_=0.07414, p-adj.=0.9967 for females and t_8_=4.144, p-adj.=0.0065 for males). **(E)** Average relative abundance of H3K18acK23ac (two-way ANOVA interaction F_1,8_=1.683, p=0.2307; sex F_1,8_=0.1001, p=0.7598; treatment F_1,8_=11.16, p=0.0102; post-hoc Šídák’s multiple comparisons t_8_=1.444, p-adj.=0.3384 for females and t_8_=3.279, p-adj.=0.0223 for males). **(F)** Average relative abundance of H2A.Z.1K4acK7ac (two-way ANOVA interaction F_1,8_=3.673, p=0.0916; sex F_1,8_=0.2963, p=0.6010; treatment F_1,8_=14.55, p=0.0051; post-hoc Šídák’s multiple comparisons t_8_=1.342, p-adj.=0.3862 for females and t_8_=4.052, p-adj.=0.0073 for males). **(G)** Venn diagrams showing overlap between genes upregulated in females with acetate (top) or downregulated in females with acetate (bottom) compared to genes upregulated in males with acetate (left, red) or downregulated in males with acetate (right, blue) in the dHPC, illustrating stark difference in response to acetate between sexes. **(H)** Heatmap of average row Z-score of TxPM for genes showing significant treatment x sex interaction (RNA-seq) during early consolidation in the dHPC. Genes showing significant interaction almost entirely corresponded to: genes upregulated in the female dHPC and downregulated in the male dHPC (n=153; shown above the dashed line) and genes downregulated in the female dHPC and upregulated in the male dHPC (n=219; shown below the dashed line). **(I)** Top 10 significantly enriched GO terms for the genes upregulated in the female dHPC and downregulated in the male dHPC in response to acetate treatment. **(J)** Top 10 significantly enriched GO terms for the DEGs downregulated in the female dHPC and upregulated in the male dHPC in response to acetate treatment. **(K)** Venn diagram showing overlap of genes with significant treatment x sex interaction in the dHPC during early consolidation and H2A.Z KO DEGs, identified by (12). For volcano plots, dots represent individual hPTMs (gray indicates no significant difference, blue significantly downregulated, and red significantly upregulated in response to acetate), numbers denote the quantity of significantly downregulated (blue, top left) or significantly upregulated (red, top right) hPTMs. For Venn diagrams, overlaps were analyzed using Fisher’s exact test (corresponding p-values and number of genes displayed). For GO terms, numbers next to bars denote the quantity of significant DEGs within each corresponding GO term. *p<0.05, **p<0.01, ***p<0.001. Sample sizes are n=3 per sex per group. Abbreviations: DEGs, differentially expressed genes; dHPC, dorsal hippocampus; GO, gene ontology; hPTMs, histone post-translational modifications; NOL, novel object location; TxPM, transcripts per million).

Surprisingly, transcriptomic analyses of pooled sexes revealed very few DEGs following acetate administration in the dHPC (11 DEGs in total; ***Fig. S4B***), while analysis of sex-stratified data resulted in considerably more DEGs (149 DEGs in females, 655 DEGs in males; ***Fig. S4C***). This may suggest sex-specific responses to acetate during consolidation, which would result in dampening of DEG detection when sexes are pooled, similar to what was observed for hPTMs in LC-MS/MS (***Fig. S2A***). In support of this, both upregulated and downregulated genes in females showed almost no overlap with upregulated and downregulated genes in males (***Fig. 6G***). To assess the degree to which females and males differed in response to acetate during consolidation, we used an interaction effect in RNA-seq analyses, testing for genes that showed a significant difference in response to acetate depending upon sex (see methods). This revealed that 372 genes showed significant interaction between sex and treatment (***Fig. S4D***), and these generally fell into two categories: 153 genes increased in females and decreased in males following acetate (***Fig. 6H, top***) and 219 genes increased in males and decreased in females (***Fig. 6H, bottom***). Gene ontology (GO) analysis revealed that genes increased in females and decreased in males were related to behavior, regulation of synaptic transmission, and neuronal action potential, suggesting an activation of gene expression programs related to neuronal activity and synaptic plasticity during the critical consolidation window (***Fig. 6I***). In contrast, genes decreased in females and increased in males were related to metabolic terms, including positive regulation of lipid biosynthesis and positive regulation of CoA-transferase activity (***Fig. 6J***). This stark transcriptional difference in response to acetate during memory consolidation could play a key role in shaping the sex-specificity of long-term behavioral outcomes.

Compared to the later time point, the relationship between acetate-regulated genes and H2A.Z-deletion sensitive genes was different during the consolidation period. In the female dHPC, acetate-upregulated genes still overlapped with genes downregulated by H2A.Z KO (Fisher’s exact test, p=3.1E-06) and acetate-downregulated genes overlapped with genes upregulated by H2A.Z KO (Fisher’s exact test, p=4.8E-15) (***Fig. S4E***). In males, these trends were opposite: acetate-upregulated genes overlapped with genes upregulated by H2A.Z KO (Fisher’s exact test, p=1.3E-21) while acetate-downregulated genes did not significantly overlap with either genes upregulated or downregulated by H2A.Z KO (***Fig. S4E***). Because of the apparent opposite relationship between sex-specific acetate changes and H2A.Z KO DEGs, we compared H2A.Z KO DEGs to the 372 genes showing significant interaction effect between treatment and sex (***Fig. 6K***). We found an even more striking pattern: over 66% of genes showing female upregulation and male downregulation with acetate overlapped H2A.Z KO downregulated genes (***Fig. 6K, light red***) and nearly 68% of genes showing female downregulation and male upregulation with acetate overlapped H2A.Z KO upregulated genes (***Fig. 6K, light blue***). These analyses, together with the mass spectrometry data, outline a close relationship between the histone variant H2A.Z and transcriptional changes, in which H2A.Z plays a distinct, sex-specific role in mediating acetate-driven gene expression during consolidation and memory recall.

Overall, we found marked differences in acetate-induced chromatin and transcriptional changes in the dHPC of female and male mice during memory consolidation, which could contribute to the sex-specific effect of acetate on long-term epigenetic and gene expression changes as well as in facilitating spatial memory formation.

## DISCUSSION

Here, we show that the intermediary metabolite acetate enhances memory formation in a sex- and brain region-specific manner. Specifically, we find that acetate facilitates dHPC-dependent spatial memory formation in female but not in male mice, while not affecting cortex-dependent non-spatial memory in either sex. This neuroactive role of the metabolite acetate is supported by previous reports indicating that acetate supplementation can regulate feeding (25) or social behavior (26). In our study, acetate-enhanced long-term memory is accompanied by chromatin changes including increased acetylation of histone variant H2A.Z, which is specific to the dHPC of female mice. Further, we find that acetate upregulates the expression of genes related to neuronal activity, synaptic plasticity and learning specifically in the female dHPC. Both early and late epigenetic and transcriptional responses to acetate differ markedly in the female and male dHPC, potentially contributing to the sex-specific behavioral outcomes.

Our findings corroborate previous reports suggesting that hippocampal histone acetylation is closely linked to improved learning and memory (3, 4). In general, histone acetylation is thought to facilitate gene expression programs that are required for the formation of long-term memories. Consequently, increasing histone acetylation by genetic or pharmacological inhibition of histone deacetylases (HDACs) enhances memory formation (4, 21, 27), while decreasing histone acetylation by overexpressing HDACs inhibits learning (28). Further, we have previously shown that acetate plays an important role in dHPC histone acetylation and spatial memory in the context of alcohol exposure (9).

Increased acetylation of H2A.Z was the most significantly induced histone post-translational modification associated with acetate-enhanced memory (***Fig. 2***). H2A.Z is a histone variant with two paralogs in vertebrates called H2A.Z.1 (also known as H2A.Z, encoded by *H2afz*) and H2A.Z.2 (also known as H2A.V, encoded by *H2afv*) (29), which differ from one another by three amino acids (30). Notably, H2A.Z has recently begun to emerge as a critical regulator of hippocampal learning and memory (11, 12, 24, 31–33), and this regulation occurs via H2A.Z-mediated gene expression changes (11, 33). For example, H2A.Z is actively evicted from chromatin in the hippocampus in response to fear conditioning, permitting gene expression required for the formation of recent and remote memories (11). Consequently, shRNA-mediated depletion of H2A.Z enhances hippocampal memory and results in learning-related transcriptional changes (32) and depleting H2A.Z by inhibiting its chaperone Tip60 promotes the consolidation of remote fear memories (24). Of note, we found that acetate treatment decreases *H2Afv* and *H2Afz* transcripts in the dHPC, further supporting the role of total H2A.Z as a memory suppressor. In contrast, others reported that brain-specific deletion of H2A.Z reduces neuronal differentiation and impairs learning and memory (31), thus while the exact role of total H2A.Z is still being elucidated, its importance in regulating dorsal hippocampal memory formation and cognitive performance is consistent across studies.

Importantly, in contrast to total H2A.Z levels, acetylation of this histone variant has been suggested to promote learning and memory. Hippocampal levels of acetylated (but not total) H2A.Z are elevated both 24 hours and 30 days after contextual fear conditioning (24), albeit further increasing H2A.Z acetylation by HDAC inhibitors does not result in additional fear memory improvements. This suggests that the outcome of acetate treatment and other interventions targeting H2A.Z acetylation might be task specific, which is supported by our current findings showing that acetate enhanced memory in NOL but not NOR. Further emphasizing the importance of H2A.Z acetylation in this context, expressing an H2A.Z acetyl-mimic (H2A.Z^KQ^) in the hippocampus facilitates memory formation, whereas expressing an acetyl-defective H2A.Z^KA^ mutant impairs learning (12).

In addition to brain region differences, we observed a second layer of specificity in that acetate only increased H2A.Z acetylation and learning-related gene expression in the dHPC of female mice, and only enhanced dHPC-dependent memory in females. This combined brain region- and sex-specificity aligns with previous reports suggesting that the effects of H2A.Z on memory are both sex- and task-specific (34, 35). For example, total H2A.Z suppresses fear memory in male but not in female mice (34). Further, deletion of both H2A.Z-encoding genes in neurons improves aversive and non-aversive memory in male mice but only improves non-aversive memory in female mice (34, 35).

Acetate-enhanced memory is characterized by marked transcriptional changes in the dHPC, which could in part be mediated by the observed histone acetylation changes including those on H2A.Z. Acetylated H2A.Z colocalizes with open chromatin regions and correlates positively with gene expression (36). Strikingly, the role of H2A.Z in neuronal activity-induced gene expression is context-specific (37) and deletion of H2A.Z results in distinct transcriptional outcomes in the hippocampus of male and female mice (12). These include, for example, decreased expression of genes required for synaptic plasticity (37). Interestingly, we found that acetate in the female (but not male) dHPC preferentially upregulates the expression of genes that are decreased by H2A.Z deletion, further emphasizing the key role of acetylated H2A.Z in orchestrating transcriptional changes that underlie acetate-enhanced memory in females.

Similarly to patterns of hPTM changes, we found that acetate preferentially affected gene expression in the dHPC compared to the cortex. For example, in the female cortex, only one gene (*Apln*, encoding for Apelin) was significantly upregulated by acetate (***Fig. 3E***). Remarkably, peptides produced from the *Apelin* precursor gene have been previously implicated as negative regulators of cortex-dependent non-spatial memory; specifically, Apelin-13 was shown to inhibit long-term memory consolidation in NOR (38). The absence of acetate-induced transcriptional and epigenetic remodeling in the female cortex, along with the upregulation of a gene that inhibits NOR, likely contributes to the task-specific behavioral outcomes observed in this study.

In contrast, acetate affected the expression of 1,793 genes in the female dHPC, and a similar dHPC bias was also evident in the male brain (***Fig. 3E***). Acetate-upregulated genes in the dHPC were enriched for genes linked to synaptic plasticity and behavior (***Fig. 3B***) and included several genes that are critical for neuronal activity and memory formation, such as *Furin*, *Plxnb2*, *Rin1* and *Wwc1*. The expression of *Furin* is reduced in Alzheimer’s disease (AD) (18), potentially contributing to impairments of dendritic spine density and memory (39). *Plxnb2* mediates hippocampal structural plasticity, and its expression is required for encoding long-term memories (17). Similarly, *Rin1* expression is required both for establishing and extinguishing fear memories (15). *Wwc1* regulates spinogenesis and cognition in mice, and polymorphisms of this gene have been linked to AD in humans (16). A key group of genes affected by acetate are response genes, which orchestrate synaptic changes required for encoding of long-term memories (40). One of the most significantly acetate-upregulated response genes in the female dHPC was *Nr4a3*, which belongs to a nuclear receptor family that has long been implicated as a key regulator of hippocampus-dependent memory formation (20). Nr4a family members regulate the expression of learning-related genes and support memory enhancement by HDAC inhibitors (21). Hippocampal *Nr4a3* expression is also increased by glucose, which is thought to be an important mechanism to enhance contextual memory formation during stress (41). The convergence of acetate and glucose on upregulating hippocampal *Nr4a3* expression emphasizes the importance of metabolic-epigenetic interactions in the brain (*1, 2*), particularly in the context of memory formation.

Notably, response genes affected by acetate were predominantly upregulated in females, while in males there was no such directional bias (***Fig. 3E***). One response gene, *Popdc3*, was significantly increased in the female dHPC and significantly decreased in the male dHPC (***Fig. 3F***). This pattern was shared by three other genes: *Nnat*, *Cpne4* and *Amigo2*. Of particular interest is *Amigo2*, which acts as a positive regulator of gene expression and synapse assembly in the hippocampus (42). Additional response genes specifically upregulated in the female dHPC included *Tgf2b*, *Fgf2*, *Fhl2*, and *Hmgcs1*. Of note, *Hmgcs1* is involved in acetyl-CoA metabolism, suggesting that the acetate – acetyl-CoA – histone acetylation pathway we and others previously described (1, 2, 9, 43) could play an important role in mediating acetate-induced chromatin and transcriptional changes in the female dHPC, which underlie female-specific behavioral outcomes.

Nevertheless, there are alternative mechanisms by which acetate could enhance learning and memory that warrant further investigation. These include decreasing the activity of HDACs (44) or serving as a precursor for glutamate and GABA synthesis (45) The functional differences between acetate-induced transcriptional remodeling in the female and male dHPC were particularly evident when we interrogated genes differentially expressed in response to NOL training (22). In females, NOL upregulated genes were significantly increased by acetate and NOL downregulated genes were significantly decreased by acetate (***Fig. 4B***), strongly suggesting that acetate remodels dorsal hippocampal gene expression programs in a manner that facilitates spatial learning. This included the increased expression of several epigenetic enzymes and response genes (***Fig. 4E***), emphasizing the critical role of chromatin and transcription-related processes in acetate-enhanced memory. Acetate- and NOL-upregulated genes in females significantly overlapped with genes that are downregulated following H2A.Z KO (***Fig. 5D***), which is in strong support of our proteomic data identifying H2A.Z acetylation as a potential regulator of acetate-induced transcriptional remodeling in the female dHPC. In stark contrast, NOL upregulated genes were significantly decreased by acetate in the male dHPC, while NOL downregulated genes were not affected by treatment in males (***Fig. 4B***). This suggests that while acetate also affects gene expression in the male dHPC, these changes are not consistent with enhanced learning, which we in fact did not observe in male mice (***Fig. 1C***).

Memory formation is a complex, time-dependent process with distinct molecular changes at different time points (20). Here, we found that the sex-specific effects of acetate on dorsal hippocampal chromatin and gene expression are present both after memory recall and early during the consolidation window (***Fig. 6***). Early hPTM changes were distinct from late hPTM changes in that they were fewer in number, more pronounced in males, and primarily affected canonical histones, including ones linked to gene activation and metabolite storage (46). In contrast, late hPTM changes were more numerous overall, more pronounced in females, and more frequently affected histone variants such as H2A.Z. Compared to memory recall, the sex-specificity of acetate-induced transcriptional changes was even more striking during early memory consolidation. In fact, female upregulated genes showed a consistent trend of downregulation in males, and female downregulated genes showed a consistent trend of upregulation in males. As “behavior” was the most enriched functional category among female-biased genes, these results suggest that early transcriptional changes in the female dHPC are critical for acetate-enhanced memory. Further underscoring the importance of epigenetic and gene expression changes during early consolidation, we have previously shown that inhibiting ACSS2 – which is required for acetate-dependent histone acetylation and gene expression changes (9) – during this time frame prevents the formation of spatial memories (47).

Overall, these results significantly advance our understanding of how intermediary metabolites regulate chromatin and gene expression in the brain, and outline the key role of acetate in facilitating learning and memory. These findings suggest that acetate exposure could be a promising, non-invasive approach to enhance memory, particularly in females, who are disproportionately affected by both age-related cognitive decline and Alzheimer’s disease (AD) (48, 49). Levels of H2A.Z accumulate in the brain during aging (32), which potentially renders the aged population especially amenable to memory enhancement via acetate-induced H2A.Z acetylation. In support, elevating acetyl-CoA levels reduces aspects of brain aging, in part by increasing histone acetylation (50). H2A.Z levels also accumulate in AD in a sex-specific manner, only affecting females both in human populations as well as in mouse models of AD (51). Importantly, we and others have recently reported that chronic dietary supplementation of acetate rescues impairments of fear memory in mouse models of AD (49, 50). Similarly, high fiber diets, which preferentially increase the levels of circulating acetate compared to other short chain fatty acids (52), were proposed to slow the progression of neurodegenerative diseases (53, 54). Here, we show that even acute exposure to acetate enhances dHPC-dependent spatial memory in females, and that these effects are mediated by sex- and brain region-specific epigenetic and transcriptional remodeling. Consequently, acetate exposure is emerging as a potential future therapeutic tool to improve learning and memory.

## MATERIALS AND METHODS

### Animals

Animal use and all experiments performed were approved by the Institutional Animal Care and Use Committee (IACUC protocol IDs 22-0418 and 23-0440). All personnel involved have been adequately trained and are qualified according to the Animal Welfare Act (AWA) and the Public Health Service (PHS) policy. C57BL/6J mice 9–10 weeks of age were used for all experiments. Mice were housed on a 12h/12h light/dark cycle, with food and water provided ad libitum.

### Behavioral assays

Behavioral experiments were conducted by the Intellectual and Developmental Disabilities Research Center (IDDRC) Animal Behavior Subunit (ABS) at Washington University in St. Louis. All behavioral experiments were conducted between 12 PM and 6 PM local time to reduce time-of-day effects. The investigator conducting behavioral assays was female and blinded to the experimental conditions. Mice were tested for the Novel Object Recognition (NOR) and Novel Object Location (NOL) tasks to assess spatial memory and associative memory, respectively. Prior to behavioral assays, mice were given a 1-week acclimation period and briefly handled for at least 3 handling sessions. Subsequently, animals went through a 2-day habituation phase, during which each mouse was habituated to the 40 x 40 cm Plexiglas testing chamber enclosed in a sound- and scent-attenuating box for 10 min on two consecutive days (***Fig. 1Ai***). For each habituation, learning, and test trial, animals were allowed to freely explore the arena for 10 min.

On Day 3, each mouse was administered either saline (vehicle control) or 1.5 g/kg acetate (75 mg/ml sodium acetate in saline solution) via intraperitoneal (i.p.) injection before the start of the NOL learning trial for object familiarization, during which two matching objects were placed in two corners of the apparatus arena and the mouse was allowed to explore for 10 min (***Fig. 1Aii, left***). On Day 4 (24 hr inter-trial interval (ITI)), each mouse was tested for the NOL task, during which one of the objects was placed in a novel location and the mouse was allowed to explore for 10 min (***Fig. 1Aii, right***). On Day 5, each mouse was again administered the appropriate treatment immediately before the start of the NOR learning trial, during which two matching objects were placed in two corners (same locations as Day 4) of the arena and the mouse was allowed to explore for 10 min (***Fig. 1Aiii, left***). On Day 6 (24 hr ITI), each mouse was tested for the NOR task, during which one of the objects was replaced with a novel object and the mouse was allowed to explore for 10 min (***Fig. 1Aiii, right***). For memory recall experiments (later timepoint, Figs 2-5), mice were sacrificed via cervical dislocation immediately upon completion of the NOL/NOR behavioral battery, which was approximately 24 hr following the final acetate exposure. For early memory consolidation experiments (early timepoint, Fig 6), mice were sacrificed via cervical dislocation immediately following the NOL training trial on Day 3, which was approximately 30 min following acetate exposure. Brains were immediately collected in a cryogenic vial and snap-frozen in a bath of 2-methylbutane (Sigma-Aldrich, Cat. No. M32631) embedded in dry ice. Whole brain tissue samples were subsequently stored at -80°C until dissection. Brain tissue was dissected on a cold plate set to -20°C, using a razor blade to create slices and a sample corer (11-gauge, Fine Science Tools Cat. No. 180-35) to collect specific brain regions.

All trials were video recorded using a camera mounted above each apparatus. ANY-Maze video tracking software (Stoelting Company, Wood Dale, Illinois, USA) was used to track animal movement (quantify distance traveled within the apparatus) and exploration of objects (quantify time spent exploring each object). Object exploration was defined as the presence of a mouse’s head within 2 cm of the object and orienting towards the object. Objects were placed centered 2 cm away from the adjacent walls of the placement corner. Objects used (3D-printed apartment or PVC and metal ring spiral) are approachable and approached with equal interest, and are not difficult to climb/stand on. Novel object and novel location were equally counterbalanced across each group to eliminate object or side bias. Between each use, the apparatus arena was cleaned with 0.02% chlorhexidine solution and the objects were cleaned with 70% ethanol to eliminate olfactory cues for the next use.

Novelty preference index (NPI) was calculated according to the following formula: (time exploring novel - time exploring familiar) / (time exploring novel + time exploring familiar))*100. Animals that did not interact with both objects during the NOL learning trial were excluded from results and were not used for further molecular analyses. All experiments were performed and analyzed by blinded investigators.

### Histone preparation and mass spectrometry

Histone extraction and sample preparation for mass spectrometry were performed as previously described (55). Brain tissue was dounce homogenized in nuclear isolation buffer (15 mM Tris-HCl, 15 mM sodium chloride, 60 mM potassium chloride, 5 mM magnesium chloride, 1 mM calcium chloride, and 250 mM sucrose at pH 7.5; with 0.5 mM AEBSF, 10 mM sodium butyrate, 5 mM microcystin, and 1 mM dithiothreitol added fresh) with 0.3 % NP-40 and incubated for 5 min on ice. Nuclei were collected by centrifugation (600 g for 5 min at 4°C). The resulting nuclear pellet was washed 2x with the same volume of nuclear isolation buffer (without NP-40). Histones were then acid-extracted with 0.2 N sulfuric acid for 4 hr at 4°C with rotation. The insoluble nuclear debris was pelleted (3,400 g for 5 min at 4°C), and the supernatant was retained. Histone proteins were then precipitated by adding 100% trichloroacetic acid in a 1:4 ratio (v/v) and incubating overnight at 4°C. The pellet was washed with acetone to remove residual acid, and histone proteins were resuspended in 30 µl of 50 nM ammonium bicarbonate (pH 8.0).

To derivatize primary and secondary amines, the histone sample was mixed with 15 µl derivatization mix (propionic anhydride and acetonitrile in a 1:3 ratio (v/v)), immediately followed by addition of 7.5 µl ammonium hydroxide to maintain pH 8.0. The sample was incubated for 15 min at room temperature, and the derivatization procedure was repeated once. Samples were then dried and resuspended in 50 mM ammonium bicarbonate and incubated with sequencing-grade modified trypsin (Promega, Cat. No. V5113) in a 1:20 enzyme:sample ratio overnight at room temperature. Following trypsin digestion, the derivatization reaction was performed twice to derivatize the N-termini of the peptides. Samples were desalted with C18 stage tips (made using Fisher Scientific, Cat. No. 13-110-019).

An LC-MS/MS system consisting of a Vanquish Neo UHPLC coupled to an Orbitrap Q Exactive or Ascend (Thermo Scientific) was used for peptide analysis. Histone peptide samples were maintained at 7°C on a sample tray in LC. Separation of peptides was carried out on an Easy-Spray™ PepMap™ Neo nano-column (2 µm, C18, 75 µm x 150 mm) at room temperature with a mobile phase. The chromatography conditions consisted of a linear gradient from 2–32% solvent B (0.1% formic acid in 100% acetonitrile) in solvent A (0.1% formic acid in water) over 48 min and then 42–98% solvent B over 12 min at a flow rate of 300 nL/min. The mass spectrometer was programmed for data-independent acquisition (DIA). One acquisition cycle consisted of a full MS scan and 35 DIA MS/MS scans of 24 m/z isolation width starting from 295 m/z to 1100 m/z. Full MS scans were typically acquired in the Orbitrap mass analyzer across 290–1200 m/z at a resolution of 70,000 or 120,000 in positive profile mode with an injection time of 50 ms and an automatic gain control (AGC) target of 1.0E-06 or 200%. MS/MS data from higher energy collisional dissociation (HCD) fragmentation was collected in the Orbitrap. These scans typically used a nominal collision energy (NCE) of 30 or 25, AGC target of 1000%, and maximum injection time of 60 ms. EpiProfile 3 (56) was used to analyze histone MS data and calculate the ratio at which each modification was present for each peptide (relative abundance).

### Western blot

Histone extracts were quantified using the Qubit^®^ Protein Assay Kit (Life Technologies, Cat. No. Q33211) with the Qubit^®^ 4 Fluorometer according to manufacturer protocols. Each Western blot (WB) sample was prepared as 3 µg histone protein in 1X NuPAGE™ LDS Sample Buffer (Invitrogen, Cat. No. NP0007), 1X NuPAGE™ Sample Reducing Agent (Invitrogen, Cat. No. NP0004), and molecular grade water. Samples were incubated for 5 min at 95°C. Wells of a 4-12% Bis-Tris Mini Protein Gel (1.0-1.5 mm, Invitrogen, Cat. No. NP0335BOX) were flushed with NuPAGE™ Antioxidant (Invitrogen, Cat. No. NP0005) and samples were then loaded onto the gel. Electrophoresis was performed at 100 V constant voltage in an XCell SureLock™ Mini-Cell apparatus (Invitrogen, Cat. No. EI0001) filled with 1X NuPAGE™ MOPS SDS Running Buffer (Invitrogen, Cat. No. NP0001). Following electrophoresis, histones were transferred onto a 2 µm nitrocellulose membrane (Bio-Rad, Cat. No. 1620112) with 25 V constant voltage for 90 min at 4°C. Following transfer, Ponceau S solution (Sigma-Aldrich, Cat. No. P7170) was used to verify even loading and successful transfer. The membrane was briefly washed 2x with Milli-Q^®^ water and imaged using an iBright™ FL1500 Imaging System (Invitrogen, Cat. No. A44241). The membrane was then placed into a dark staining box, briefly washed with 1X TBST (EZ BioResearch, Cat. No. S-1012), and blocked in blocking solution (5% milk in 1X TBST) for 1 hr at room temperature with agitation. The membrane was incubated in 1:1000 H2A.Z pan-acetyl (K4/K7/K11/K13) polyclonal antibody (Invitrogen, Cat. No. PA5-114690, Lot No. ZD4298112, RRID AB_2899326; diluted in blocking solution) overnight at 4°C with agitation. The next day, the membrane was washed in 1X TBST 3x 10 min each at room temperature with agitation, was incubated in 1:1000 HRP secondary antibody (Invitrogen, Cat. No. A27036, Lot No. 2567325, RRID AB_2536099; diluted in blocking solution) for 1 hr at room temperature with agitation and was again washed in 1X TBST 3x 10 min each at room temperature with agitation. The membrane was incubated in SuperSignal™ West Pico PLUS Chemiluminescent Substrate (Thermo Scientific, Cat. No. 34577) for 5 min at room temperature with agitation prior to imaging using an iBright™ FL1500 Imaging System.

### RNA extraction and sequencing

Total RNA was extracted using TRIzol-chloroform and the RNeasy Mini Kit (Qiagen, Cat. No 74104) including RNase-free DNase (Qiagen, Cat. No. 79254) treatment according to manufacturer protocols. Total RNA quality was assessed using High Sensitivity RNA ScreenTape Analysis on the TapeStation system (Agilent, Cat. No 5067-5579 and 5067-5580) according to manufacturer protocols.

mRNA was isolated from up to 150 ng total RNA using the NEBNext® Poly(A) mRNA Magnetic Isolation Module (New England Biolabs, Cat. No. E7490), and libraries were prepared using the NEBNext® Ultra™ II RNA Library Prep Kit for Illumina® (New England Biolabs, Cat. No. E7770) using compatible NEBNext® Oligos for Illumina (New England Biolabs, Cat. No. E7335 and E7500). Library quality was assessed using D1000 DNA ScreenTape Analysis on the TapeStation system (Agilent, Cat. No. 5067-5602 and 5067-5582). Libraries were quantified using the NEBNext® Library Quant Kit (New England Biolabs, Cat. No. E7630) on an Applied Biosystems™ QuantStudio™ 5 Real-Time PCR System. The NEBioCalculator qPCR Library Quantification Tool was used to calculate the concentration of amplifiable templates in each library. Following quantification, libraries were pooled and submitted for sequencing.

Libraries were sequenced on the AVITI system using the AVITI 2×75 Sequencing Kit Cloudbreak Freestyle (FS) High Output Kit (Element Biosciences, Cat. No. 860-00015) by the DNA Sequencing Innovation Lab (DSIL) at The Center for Genome Sciences and Systems Biology (CGSSB) at Washington University School of Medicine.

### RNA-seq analysis

Adapters were trimmed from reads using Cutadapt V3.5 (57) with a minimum length of 15 bp. Reads were aligned to the *Mus musculus* GRCm39 reference genome assembly using STAR v2.7.10a (58). STAR alignments were performed in two passes, with un-annotated splice junctions featuring >2 unique reads per replicate were incorporated into the reference followed by remapping of the libraries. Gene-level counts were generated using featureCounts, using the RefSeq annotation (GCF_000001635.27) containing all protein coding and long noncoding RNAs. Differential gene expression analyses were performed with DESeq2 (59). For all pairwise comparisons, the Wald negative binomial test (test=“Wald”) was used for determining DEGs. Unless otherwise stated, an adjusted p-value cutoff of 0.1 was used in differentiating differentially expressed from non-differing genes to maximize the sensitivity of RNA-seq results. Gene Ontology (GO) enrichment tests were performed with the R package topGO (60), using the fishers elim method. For gene overlap calculations Fisher’s exact tests were used, ensuring that the total ‘universe’ of genes tested represented an intersection of each sets *testable* genes, omitting those with insufficient data (DESeq2 padj=”NA”).

For sex-specific responses to acetate (e.g., ***Figs. 4B,D, 5B, S4E***) a given sex’s acetate samples were compared to all others for that tissue after blocking for sex. For the interaction effect (***Figs. 6H-K, S4D***) the full model: ∼sex+treatment+sex:treatment was compared to the reduced model ∼sex+treatment, utilizing a likelihood ratio test (test=”LRT” in DESeq2) to identify genes whose changes with acetate differ significantly depending on treatment.

Volcano plots were generated using enhancedvolcano (61). For heatmaps, row Z-scores were calculated on averaged transcripts per million (TxPM). Heatmaps were generated using heatmap.2 in the gplots R package (62), which uses Euclidean measures to obtain distance matrices and complete agglomeration for clustering. MA plots were generated using ggplot2 (62).

### Statistical analysis of behavioral and LC-MS/MS data

Sample sizes and statistical tests are indicated in figure legends. Statistical tests were performed in GraphPad Prism with significance threshold *p<0.05 unless otherwise stated. Bar plots present mean ± SEM. Boxplots and violin plots present standard interquartile ranges and median. Shapes represent individual mice (gray circles saline-treated and green squares acetate-treated) in bar plots, boxplots, and violin plots.

For behavior data, data distribution was tested using the Shapiro-Wilk normality test. Outliers, if present, were detected using Grubbs’ test and were excluded from analyses. Parametric data were analyzed using Student’s t test (unpaired, two-tailed, equal variance; acetate vs. saline, sexes combined) or two-way ANOVA (data stratified by sex) to detect significant interactions, followed by post-hoc Šídák’s multiple comparisons test to detect specific significant differences. Nonparametric data were analyzed using Mann-Whitney U test (acetate vs. saline, data for sexes pooled) or multiple Mann-Whitney tests followed by Holm-Šídák correction for multiple comparisons (data stratified by sex). Mice that did not interact with both objects during the NOL learning trial were excluded from analysis.

Initial statistical analyses for LC-MS/MS data were performed using Student’s t test (unpaired, two-tailed, equal variance) for each histone peptide relative abundance ratio (per row) to compare acetate-vs. saline-treated groups. For hPTMs of interest, relative abundance data were further analyzed using two-way ANOVA (data stratified by sex) to detect significant interactions, followed by Šídák’s multiple comparisons test to detect specific significant differences.

## ACKNOWLEDGEMENTS

We thank Katie McCullough and Susan Maloney for their invaluable help in performing the behavioral analyses. This work was supported by NIH grants R00AA028577 (GE), Alzheimer’s Association grant AARF-19-618159 (GE), 7R01AI118891 (BAG), R01HD106051 (BAG), a Small Grant from the McDonnell Center for Cellular and Molecular Neurobiology (GE) and the NARSAD Young Investigator Grant YIG31527 from the Brain and Behavior Research Foundation (GE). Behavioral research reported in this publication was supported by the Eunice Kennedy Shriver National Institute of Child Health and Human Development of the National Institutes of Health under Award Number P50 HD103525 to the Intellectual and Developmental Disabilities

Research Center (IDDRC), as well as by funds provided by the McDonnell Center for Systems Neuroscience, McDonnell Center for Cellular and Molecular Neuroscience, and the Taylor Family Institute at Washington University in St. Louis. RNA sequencing was conducted by the DNA Sequencing Innovation Lab (DSIL) at The Center for Genome Sciences and Systems Biology (CGSSB) at Washington University School of Medicine in St. Louis.

## Author contributions

Conceptualization: GE; Data curation: EMP, KMG, GE; Formal analysis: EMP, KMG, FNV, GE; Funding acquisition: GE; Investigation: EMP, KMD, FNV, GE; Methodology: GE, KMG; Project administration: GE; Resources: KMG, FNV, BAG, GE; Software: KMG; Supervision: GE; Validation: EMP, GE; Visualization: EMP, KMG; Writing – original draft: EMP, GE.

## Data availability

All mass spectrometry and sequencing data will be made publicly available at the time of publication.

---

**Supplementary Figure S1.**
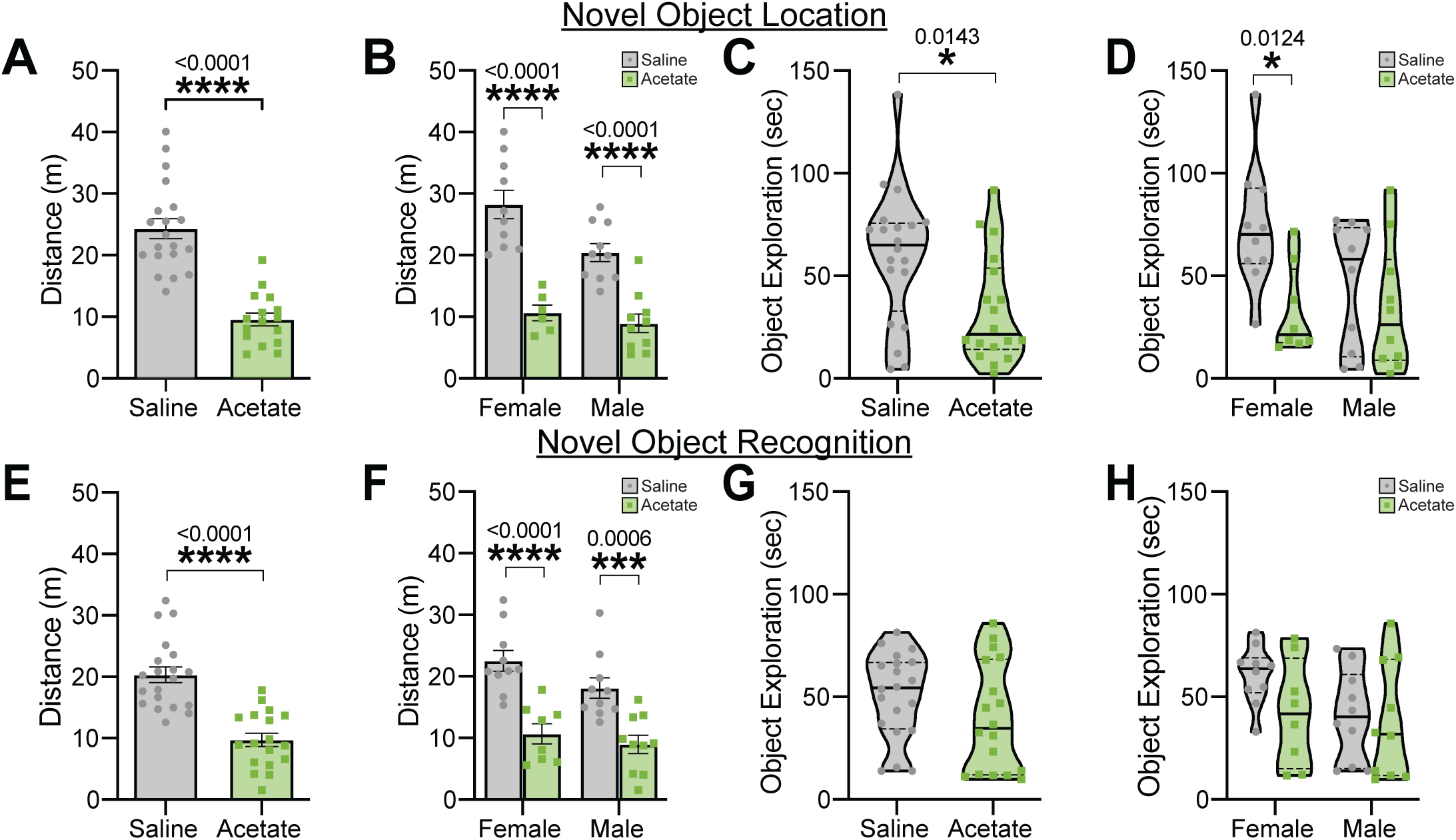
Acetate induces acute hypoactivity in mice. **(A-D)** Distance traveled (m) and object exploration time (sec) during the NOL learning trial. **(A)** Average distance traveled (m) during the NOL learning trial (Student’s t test t_34_=7.269, p<0.0001). **(B)** Average distance traveled (m) during the NOL learning trial, grouped by sex (two-way ANOVA interaction F_1,32_=2.797, p=0.1042; sex F_1,32_=6.724, p=0.0142; treatment F_1,32_=62.58, p<0.0001; post-hoc Šídák’s multiple comparisons t_32_=6.339, p-adj.<0.0001 for females and t_32_=4.765, p-adj.<0.0001 for males). **(C)** Object exploration time (sec) during the NOL learning trial (Mann-Whitney U=97, p=0.0143). **(D)** Object exploration time (sec) during the NOL learning trial, grouped by sex (multiple Mann-Whitney U=10, p=0.006216 for females and U=39, p=0.4241 for males; post-hoc Holm-Šídák multiple comparisons correction p-adj.=0.0124 for females and p-adj.=0.4241 for males). **(E-H)** Distance traveled (m) and object exploration time (sec) during the NOR learning trial. **(E)** Average distance traveled (m) during the NOR learning trial (Student’s t test t_34_=8.575, p<0.0001). **(F)** Average distance traveled (m) during the NOR learning trial, grouped by sex (two-way ANOVA interaction F_1,34_=0.6859, p=0.4133; sex F_1,34_=3.458, p=0.0716; treatment F_1,34_=40.82, p<0.0001; post-hoc Šídák’s multiple comparisons t_34_=4.960, p-adj.<0.0001 for females and t_34_=4.053, p-adj.=0.0006 for males). **(G)** Object exploration time (sec) during the NOR learning trial (Mann-Whitney U=129, p=0.1392). **(H)** Object exploration time (sec) during the NOR learning trial, grouped by sex (multiple Mann-Whitney U=22, p=0.1220 for females and U=40, p=0.4686 for males; post-hoc Holm-Šídák multiple comparisons correction p-adj.=0.2291 for females and p-adj.=0.4686 for males). *p<0.05, ***p<0.001, ****p<0.0001. For distance, sample sizes are n_Saline,F_=10, n_Saline,M_=10, n_Acetate,F_=6, n_Acetate,M_=10. Object exploration data are non-parametric and sample sizes are n_Saline,F_=10, n_Saline,M_=10, n_Acetate,F_=8, n_Acetate,M_=10. Abbreviations: m, meters; NOL, novel object location; NOR, novel object location; sec, seconds.

**Supplementary Figure S2.**
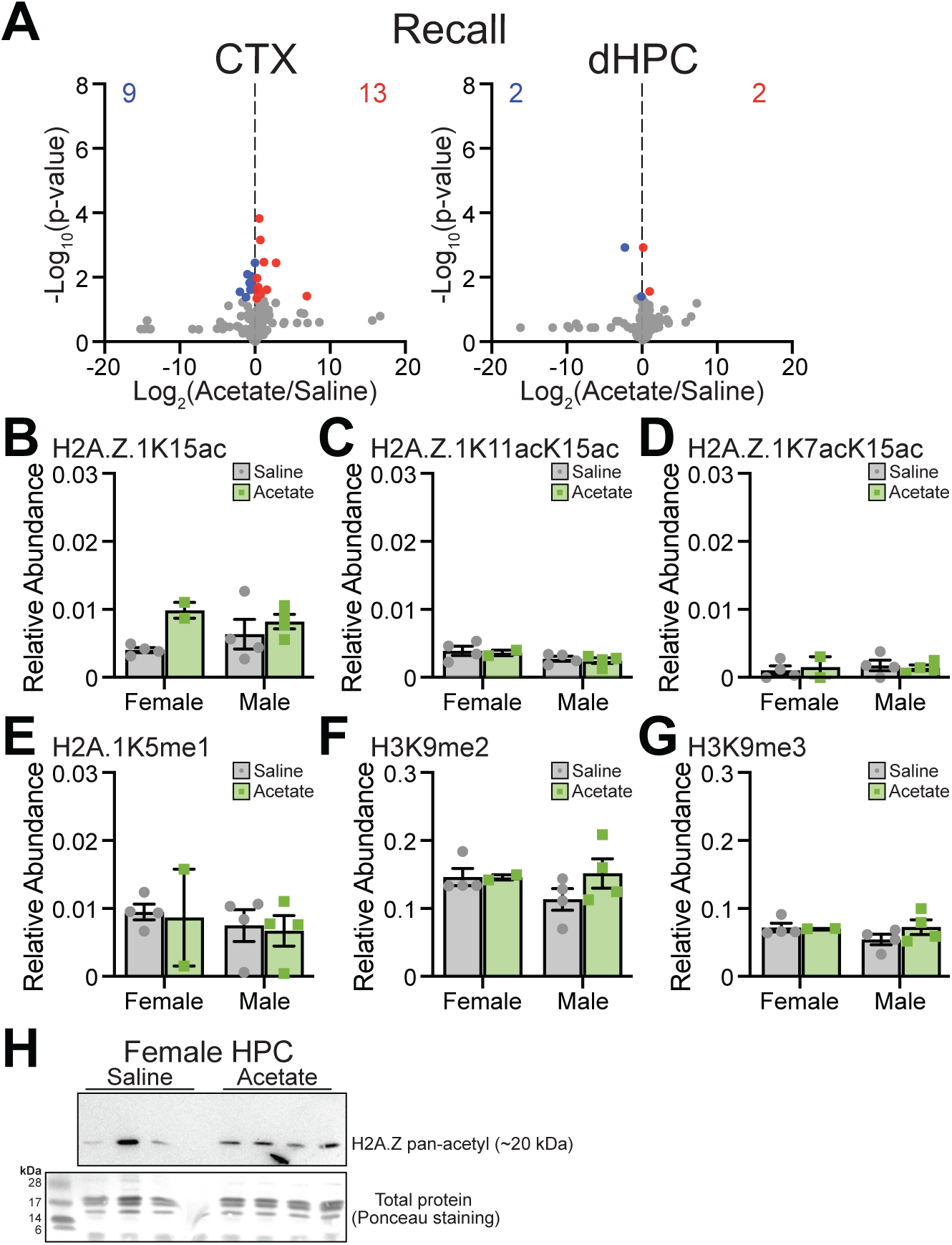
Effects of acetate on histone modifications following memory recall. **(A)** Volcano plots of hPTMs in the CTX (left) and dHPC (right) comparing acetate- and saline-treated mice (both sexes) sacrificed following memory recall (late timepoint). **(B-G)** Average relative abundance of hPTMs of interest in the CTX following memory recall. **(B)** Average relative abundance of H2A.Z.1K15ac (two-way ANOVA interaction F_1,10_=1.682, p=0.2238; sex F_1,10_=0.04827, p=0.8305; treatment F_1,10_=6.264, p-adj.=0.0313; post-hoc Šídák’s multiple comparisons t_10_=2.453, p=0.0670 for females and t_10_=0.9535, p-adj.=0.5970 for males). **(C)** Average relative abundance of H2A.Z.1K11acK15ac (two-way ANOVA interaction F_1,10_=2.61E^-04^, p=0.9874; sex F_1,10_=4.445, p=0.0612; treatment F_1,10_=0.2736, p=0.6123; post-hoc Šídák’s multiple comparisons t_10_=0.3480, p-adj.=0.9298 for females and t_10_=0.4007, p-adj.=0.9082 for males). **(D)** Average relative abundance of H2A.Z.1K7acK15ac (two-way ANOVA interaction F_1,10_=0.2738, p=0.6122; sex F_1,10_=0.1610, p=0.6967; treatment F_1,10_=4.91E-03, p=0.9455; post-hoc Šídák’s multiple comparisons t_10_=0.3830, p-adj.=0.9158 for females and t_10_=0.3583, p-adj.=0.9258 for males). **(E)** Average relative abundance of H2A.1K5me1 (two-way ANOVA interaction F_1,10_=5.12E-05, p=0.9944; sex F_1,10_=0.5138, p=0.4899; treatment F_1,10_=0.08544, p=0.7760; post-hoc Šídák’s multiple comparisons t_10_=0.1933, p-adj.=0.9777 for females and t_10_=0.2254, p-adj.=0.9698 for males). **(F)** Average relative abundance of H3K9me2 (two-way ANOVA interaction F_1,10_=1.138, p=0.3112; sex F_1,10_=0.5542, p=0.4737; treatment F_1,10_=1.073, p=0.3247; post-hoc Šídák’s multiple comparisons t_10_=0.01992, p-adj.=0.9998 for females and t_10_=1.662, p-adj.=0.2386 for males). **(G)** Average relative abundance of H3K9me3 (two-way ANOVA interaction F_1,10_=1.169, p=0.3050; sex F_1,10_=0.7247, p=0.4145; treatment F_1,10_=0.8592, p=0.3758; post-hoc Šídák’s multiple comparisons t_10_=0.09964, p-adj.=0.9940 for females and t_10_=1.588, p-adj.=0.2663 for males). **(H)** Western blot of pan-acetylated H2A.Z in the hippocampus (HPC) of female mice 24 hours following i.p. injection of 1.5 g/kg acetate (sample sizes are n=4/treatment). Ponceau stain is shown for total protein content. For volcano plots, dots represent individual hPTMs (gray not significant, blue significantly downregulated, and red significantly upregulated by acetate), numbers denote the quantity of significantly downregulated (blue, top left) or significantly upregulated (red, top right) hPTMs. *p<0.05, **p<0.01, ***p<0.001. Sample sizes are n=3 per sex per group unless otherwise specified. Abbreviations: dHPC, dorsal hippocampus; HPC, hippocampus; hPTMs, histone post-translational modifications; NOL, novel object location.

**Supplementary Figure S3.**
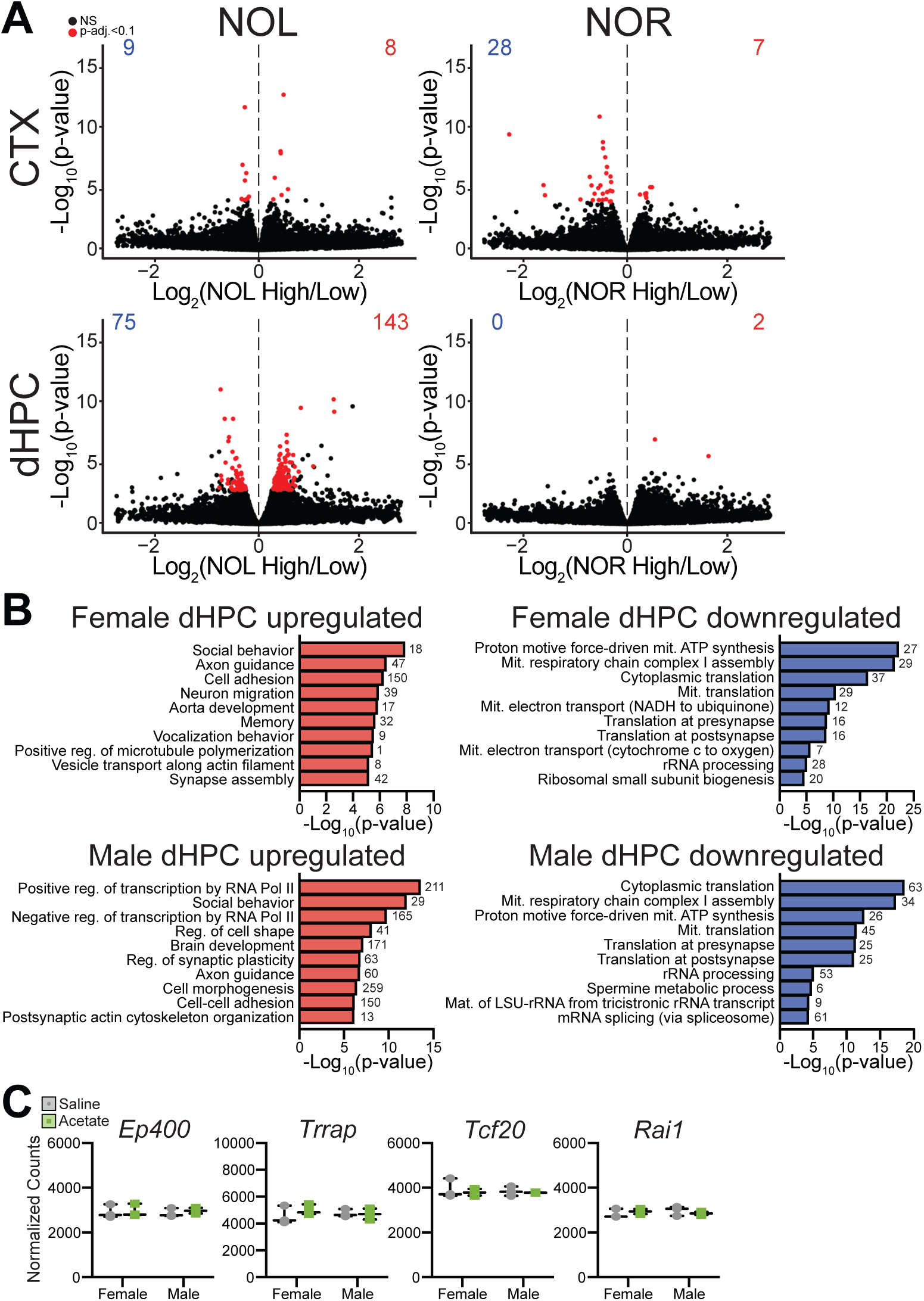
Effects of acetate on gene expression following memory recall. **(A)** Volcano plots showing differential gene expression in the CTX (top) and dHPC (bottom) for learners vs. non-learners in NOL (left) or NOR (right). Learners are characterized by a positive NPI for the respective test, whereas non-learners have a negative NPI for the respective test. Dots represent individual genes (black not significant (NS), red significant (p-adj.<0.1)) and numbers denote the quantity of significantly downregulated (blue, top left) or significantly upregulated (red, top right) genes (learners vs. non-learners). **(B)** Top 10 significantly enriched GO terms in the dHPC of females (top) and males (bottom) among genes upregulated (left) and downregulated (right) by acetate following memory recall. Numbers next to bars denote the quantity of significant DEGs within each corresponding enriched GO term. **(C)** Normalized gene counts in the CTX for GOIs associated with H2A.Z, identified by (23). *Ep400* and *Trrap* encode components of the TIP60/p400 complex, and *Tcf20* and *Rai1* encode components of the PHF14 complex. Sample sizes are n=3 per sex per group. Abbreviations: CTX, cortex; DEGs, differentially expressed genes; dHPC, dorsal hippocampus; Ep400, E1A binding protein p400; GO, gene ontology; GOIs, genes of interest; mat., maturation; mit., mitochondrial; NOL, novel object location; NOR, novel object recognition; NPI, novelty preference index; NS, not significant; PHF14, PHD finger protein 14; Rai1, retinoic acid induced 1; reg., regulation; Tcf20, transcription factor 20; TIP60 (KAT5), lysine acetyltransferase 5; Trrap, transformation/transcription domain-associated protein.

**Supplementary Figure S4.**
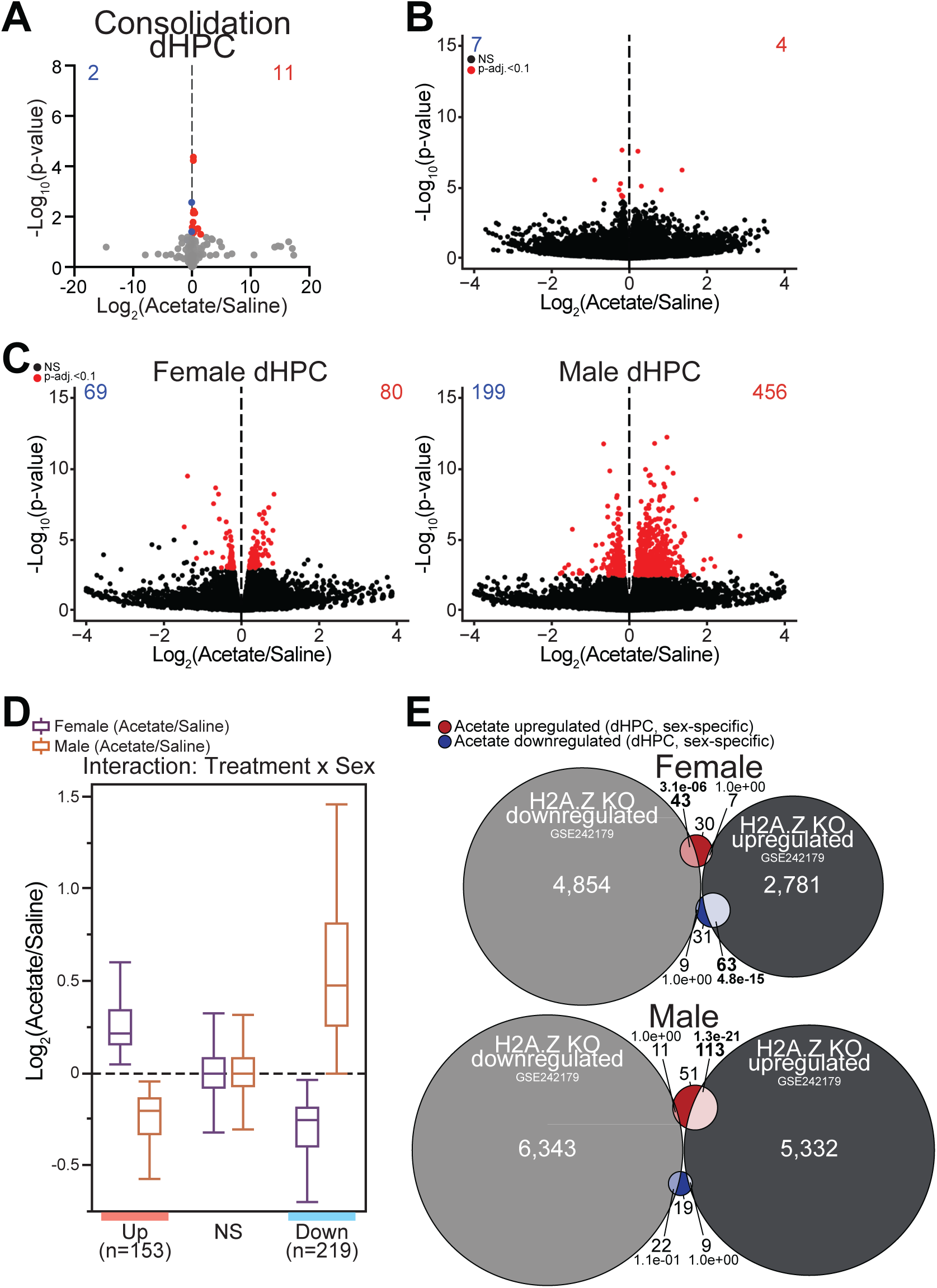
Effects of acetate on histone modifications and gene expression during consolidation. **(A)** Volcano plot of hPTMs in the dHPC comparing acetate- and saline-treated mice (both sexes) sacrificed following the NOL learning trial, during consolidation (early timepoint). **(B)** Volcano plot showing differential gene expression in the dHPC comparing acetate- and saline-treated mice sacrificed following the NOL learning trial, during consolidation (early timepoint). **(C)** Volcano plots showing differential gene expression in the dHPC comparing acetate- and saline-treated females (left) and males (right) sacrificed during consolidation. **(D)** Boxplots showing log2(fold change) in females (blue) and males (red) for treatment x sex interaction DEGs during consolidation, illustrating that interaction DEGs largely show opposite regulation upon acetate treatment in the two sexes. **(E)** Venn diagrams showing overlap of DEGs in response to acetate treatment (acetate upregulated red, acetate downregulated blue) during early consolidation and H2A.Z-KO DEGs, identified by (12) in the dHPC of females (top) and males (bottom). For volcano plots, dots represent individual hPTMs (gray not significant, blue significantly downregulated, and red significantly upregulated by acetate) or transcripts (black not significant (NS), red significant (p-adj.<0.1)) and numbers denote the quantity of significantly downregulated (blue, top left) or significantly upregulated (red, top right) genes (acetate vs. saline). Sample sizes are n=3 per sex per group. Abbreviations: DEGs, differentially expressed genes; dHPC, dorsal hippocampus; KO, knockout.

